# A taxonomic synopsis of unifoliolate continental African *Vepris* (Rutaceae) with three new threatened forest tree species from Kenya and Tanzania

**DOI:** 10.1101/2022.07.16.500287

**Authors:** Martin Cheek, W.R. Quentin Luke

## Abstract

Descriptions and illustrations are presented for three new species to science, *Vepris udzungwa* Cheek*, V. lukei* Cheek (both Udzungwa Mts, Tanzania) and *V. robertsoniae* Cheek & Q. Luke (SE Kenyan kaya forests) in the context of a synoptic taxonomic revision, with an identification key to all the known unifoliolate taxa of *Vepris* in continental Africa. The remaining species are given skeletal taxonomic treatments (lacking descriptions). One widespread species in montane eastern Africa is renamed as *Vepris simplex* Cheek because its previous name, *Vepris simplicifolia* (Engl.)Mziray is predated by *Vepris simplicifolia* Endl. (basionym of *Sarcomelicope simplicifolia* (Endl.)T.G. Hartley, a widespread species of Australia, Lord Howe and Norfolk Islands, and of New Caledonia).

Conservation assessments are presented for all species, or provisional conservation assessments are presented. Of the 13 taxa, 11 are considered threatened, of which six are VU, two EN and three CR, of which two are possibly extinct globally in the Uluguru Mts of Tanzania although not yet Red Listed on iucnredlist.org.

## Introduction

Three new *Vepris* species are described in the context of a synoptic treatment of the African unifoliolate species. The research was supported by preparation for a taxonomic revision of African *Vepris* by the first author, and of floristic work for conservation prioritisation in the surviving forests of Kenya and Tanzania by the second author. The paper builds on the foundation laid for the three western African unifoliolate species by Lachenaud & Onana (2021), and increases the number of described unifoliolate *Vepris* species for continental Africa from 10 to 13. Unifoliolate *Vepris* species are likely not a natural group, but may have arisen more than once from ancestral trifoliolate species. Yet without a well-sampled phylogeny it is difficult to be certain.

*Vepris* Comm. ex A. Juss. (Rutaceae-Toddalieae), is a genus with 93 accepted species, 23 in Madagascar and the Comores and 69 in Continental Africa with one species extending to Arabia and another endemic to India (Plants of the World Online, continuously updated). The genus was last revised for tropical Africa by Verdoorn (1926). Founded on the Flore du Cameroun account of Letouzey (1963), nine new species were recently described from Cameroon (Onana & Chevillotte 2015; Cheek *et al*. 2018a; Onana *et al*. 2019; Cheek & Onana 2021; Cheek *et al*. 2022a), taking the total in Cameroon to 24 species, the highest number for any country globally, followed by Tanzania (16 species). The greatest concentrations of *Vepris* species in Cameroon are within the Cross-Sanaga Interval (Cheek *et al*. 2001) with 15 species of *Vepris* of which nine are endemic to the Interval. The Cross-Sanaga has the highest species and generic diversity per degree square in tropical Africa including endemic genera such as *Medusandra* Brenan (Peridiscaceae, Breteler *et al*. 2015; Soltis *et al*. 2007; Barthlott *et al*. 1996; Dalgallier *et al*. 2020). Much of this diversity is associated with the Cameroon Highland areas, different highlands each having a species of a genus e.g. as in *Kupeantha* Cheek (Rubiaceae, Cheek *et al*. 2018b). By comparison, neighbouring Gabon has just seven species of *Vepris* (Sosef *et al*. 2006) and just one species, *Vepris lecomteana* (Pierre) Cheek & T. Heller is listed for Congo-Brazzaville (Plants of the World Online, continuously updated).

In continental Africa, *Vepris* are easily recognised. They differ from all other Rutaceae because they have digitately (1 –)3(– 5)-foliolate (not pinnate) leaves, and unarmed (not spiny) stems. The genus consists of evergreen shrubs and trees, predominantly of tropical lowland evergreen forest, but with some species extending into submontane forests and some into drier forests and woodland. *Vepris* species are often indicators of good quality, relatively undisturbed evergreen forest since they are not pioneers.

Species of *Vepris* in Africa extend from South Africa, e.g. *Vepris natalensis* (Sond.) Mziray, to the Guinean woodland in the fringes of the Sahara Desert *(Vepris heterophylla* (Engl.) Letouzey). Mziray (1992) subsumed the genera *Araliopsis* Engl.*, Diphasia* Pierre*, Diphasiopsis* Mendonça, *Oricia* Pierre, *Oriciopsis* Engl., *Teclea* Delile, and *Toddaliopsis* Engl. into *Vepris*, although several species were only formally transferred subsequently (e.g. Harris 2000; Gereau 2001; Cheek *et al*. 2009; Onana & Chevillotte 2015). Mziray’s conclusions were largely confirmed by the molecular phylogenetic studies of Morton (2017) but Morton’s sampling was limited, identifications appeared problematic (several species appear simultaneously in different parts of the phylogenetic trees) and more molecular work would be desirable. Morton studied about 14 taxa of *Vepris*, all from eastern Africa. More recently Appelhans & Wen (2020) focussing on Rutaceae of Madagascar have found that the genus *Ivodea* Capuron is sister to *Vepris* and that a Malagasy *Vepris* is sister to those of Africa. However, the vast majority of the African species including all those of West and Congolian Africa, remain unsampled leaving the possibility open for changes to the topology of the phylogenetic tree when this is addressed.

Characteristics of some of the formerly recognised genera are useful today in grouping species. The “araliopsoid” species have hard, non-fleshy, subglobose, 4-locular fruit with 4 external grooves; the “oriciopsoid” soft, fleshy 4-locular syncarpous fruit; “oricioid” species are 4-locular and apocarpous in fruit; the fruits of “diphasioid” species are laterally compressed in one plane, bilocular and bilobed at the apex; while “tecleoid” species are unilocular in fruit and 1-seeded, lacking external lobes or grooves. There is limited support for these groupings in Morton’s study,

Due to the essential oils distributed in their leaves, and the alkaloids and terpenoids distributed in their roots, bark and leaves, several species of *Vepris* have traditional medicinal value (Burkill 1997). Burkill details the uses, essential oils and alkaloids known from five species in west Africa: *Vepris hiernii* Gereau (as *Diphasia klaineana* Pierre), *Vepris suaveolens* (Engl.) Mziray (as *Teclea suaveolens* Engl.), *Vepris afzelii* (Engl.) Mziray (as *Teclea afzelii* Engl.), *Vepris heterophylla* (Engl.) Letouzey (as *Teclea sudanica* A. Chev.) and *Vepris verdoorniana* (Exell & Mendonça) Mziray (as *Teclea verdoorniana* Exell & Mendonça) (Burkill 1997: 651 – 653). Research into the characterisation and anti-microbial and anti-malarial applications of alkaloid and limonoid compounds in *Vepris* is active and ongoing (e.g., Atangana *et al*. 2017), although sometimes published under generic names no longer in current use, e.g. Wansi *et al*. (2008). Applications include as synergists for insecticides (Langat 2011). Cheplogoi *et al*. (2008) and Imbenzi *et al*. (2014) respectively list 14 and 15 species of *Vepris* that have been studied for such compounds. A review of ethnomedicinal uses, phytochemistry, and pharmacology of the genus *Vepris* was recently published by Ombito *et al*. (2020), listing 213 different secondary compounds, mainly alkaloids and furo- and pyroquinolines, isolated from 32 species of the genus, although the identification of several of the species listed needs checking. However, few of these compounds have been screened for any of their potential applications. Recently, Langat *et al*. (2021) have published three new acridones and reported multi-layered synergistic anti-microbial activity from *Vepris gossweileri* (I.Verd.)Mziray, recently renamed as *Vepris africana* (Hook.f ex Benth.) Lachenaud & Onana (Lachenaud & Onana 2021). There is no doubt that new compounds will continue to be discovered as chemical investigation of *Vepris* species continues.

## Materials and Methods

This taxonomic study is based on herbarium specimens predominantly at EA, BM and K, field observations in Guinea, and Republic of Congo by the first author and field observations of live material in Kenya and Tanzania by the second author. All specimens seen are indicated “!”. The specimens were mainly collected using the patrol method as indicated e.g. in Cheek & Cable (1997). Herbarium citations follow Index Herbariorum (Thiers *et al*. continuously updated), nomenclature follows Turland *et al*. (2018) and binomial authorities follow IPNI (continuously updated). Material of the new species was compared morphologically with material of all other African *Vepris*, principally at K, but also using material and images from BM, EA, BR, FHO, G, GC, HNG, P and YA. Herbarium material was examined with a Leica Wild M8 dissecting binocular microscope fitted with an eyepiece graticule measuring in units of 0.025 mm at maximum magnification. The drawing was made with the same equipment using a Leica 308700 camera lucida attachment. The description was made following the format of Cheek *et al*. (2022) using terms from Beentje & Cheek (2003). Specimen location data is given as on the label of the specimens, understanding that the units formerly termed “Districts” in Kenya and Tanzania are currently termed Counties.

For the extinction risk assessment, points were georeferenced using locality information from herbarium specimens.). The conservation assessment was made using the categories and criteria of IUCN (2012), EOO was calculated with GeoCat (Bachman *et al*. 2011).

### Taxonomic Results

#### KEY TO THE UNIFOLIOLATE AFRICAN SPECIES OF *VEPRIS*

1.Leaves opposite at apex of stem; leaflets not articulated with petiole; fruit 4-loculed. W. Africa from Guinea-Conakry to Liberia

1.Leaves always alternate; leaflets articulated with petiole; fruit 1- or 2-loculed. Central to eastern Africa

2.Stems hairy (visible at stem apex with x10 hand-lense)

2.Stems glabrous (not visible at stem apex with hand-lense)

3. Stems with hairs dense; stems sparsely lenticellate (<20% cover); leaf apex acute or acuminate; petiole mostly >0.6 cm long

3. Stems with hairs sparse, erect, stems densely (>50% in patches) lenticellate, leaf apex rounded; petiole 0.3 – 0.6 cm long. Tanzania, Udzungwa Mts

4. Stems minutely puberulous; petiole (0.3 –)1 cm long; lateral nerves 20+ on each side of the midrib; Mozambique

4. Stems densely long-hairy; petioles 1.25 – 2.5 cm long; lateral nerves c. 10 – 15 on each side of the midrib. Uluguru Mts Tanzania

5. Petioles winged. Tanzania, Udzungwa Mts

5. Petioles canaliculate to cylindric. Ethiopia to Angola

6. Leaves smelling of bad fish when live (crushed) or dried; petiole 0.5 – 1.8(– 2.8) cm long; <300 m. alt. S.E. Kenya

6. Leaves smelling of *Citrus* when live (crushed) or lacking scent; petiole mostly >(1.5 –) 3 cm long (except *V. eugeniifolia* and *V. amaniensis* in E Africa, and (W. Africa) *V. africana*)>300 m alt

7. Inflorescence glabrous; stamens 8 in male flowers (4 – 7 in *V. amaniensis*)

7. Inflorescence hairy (hairs often minute); stamens 4 in male flowers

8. Fruits when ripe black; northern Angola

8. Fruits orange or red when ripe; E. Africa

9. Leaves ovate, 3.5 – 9 cm long. Somalia to Tanzania

9. Leaves elliptic, 11 – 29 cm long. Tanzania

10. Leaves leathery; petiole terete at apex; inflorescence paniculate, 9 cm long, few-flowered; stamens about twice as long as petals. Tanzania, Uluguru Mts

10. Leaves papery; petiole canaliculate at apex; inflorescence racemose 0.9 – 4(– 5) cm long; stamens shorter than petals. Tanzania, Usambara Mts

11. Lateral nerves 16 – 23 on each side of the midrib. S. Tomé, Gabon-Angola

11. Lateral nerves <14 on each side of the midrib. E Africa

12. Fruit asymmetric at base; pedicel 1 – 6 mm long. Kenya

12. Fruit symmetrical; pedicel mostly <1 mm long. Ethiopia, Kenya, Tanzania

##### 1. Vepris laurifolia

(Hutch. & Dalziel) O. Lachenaud (Lachenaud & Onana 2021: 112). Type: Guinea, Ninia, Talla Hills, 17 Feb. 1892, *Scott-Elliott* 4086 (Holotype BM barcode BM000798360!). (Fig. 1)

**Fig. 1.**
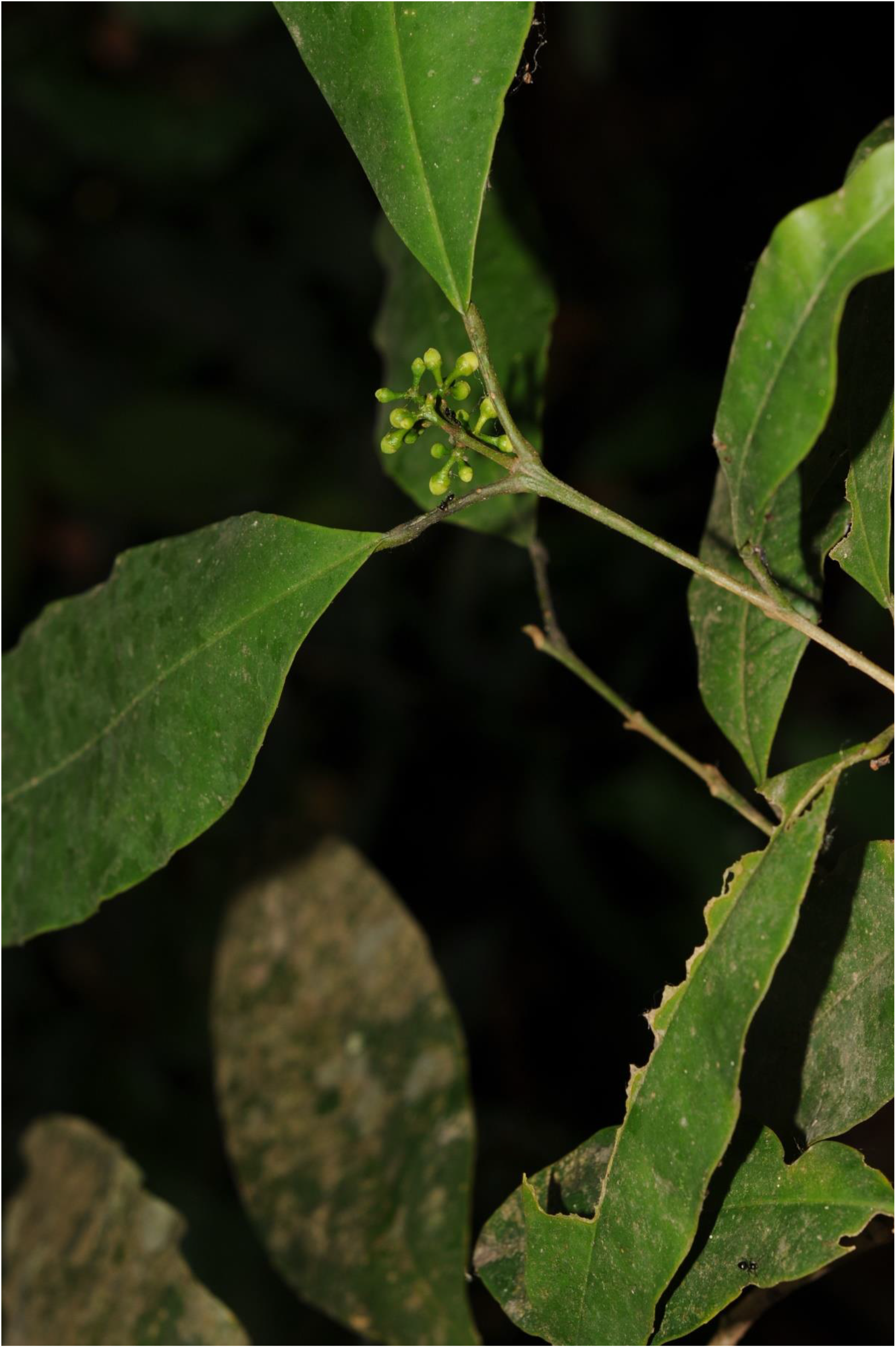
Vepris laurifolia. Photo showing habit of flowering plant (*Cheek* 16600, HNG, K) in habitat near Madina Oula, Republic of Guinea near the border with Sierra Leone, in 2012. Photo by M. Cheek.

*Garcinia laurifolia* Hutch. & Dalziel (1927: 236) *Vepris felicis* Breteler (1995: 131; Hawthorne & Jongkind 2006: 704). Type: Liberia, Central Province, c. 5 km SE of Zuole fl. 2 Apr. 1962, *J.J.F.E. de Wilde & Voorhoeve* 3754 (Holotype WAG; isotypes A, B, BR, K barcode K000800952!, P)

###### DISTRIBUTION

Guinea-Conakry, Sierra Leone, Liberia, Ivory Coast

###### REPRESENTATIVE SPECIMENS EXAMINED

**Guinea-Conakry. Guinée-Maritime**. Frigiya village, about 20 km NE of Madina Oula, fl. 29 Apr. 2012, *Cheek et al*. 16600 (HNG!, K!); After Kouria (on Coyah to Kindia road), beyond town at foot of table mts, along valley and up stream to Forest Patch 20 at head, 1 Oct. 2015, st. *Cheek* 18224 (HNG!, K!); **Guinée Forestiere,** Seredou Village, fl. 14 Feb. 2014, *P.K. Haba* (HNG, K!, WAG); Yomou prefecture. A Tayiébah, au village Kilikpala, Forêt Classée de Diécké, st. 15 Sept. 2015, *P.M. Haba* 899 (HNG, K !).

###### HABITAT

This shrub is known from lowland evergreen forest, usually associated with water courses (possibly because other areas have been cleared). The altitudinal range is 80 – 624 m. In the field notes of several specimens it is described as being found in ‘forest relics’ suggesting that it only occurs in patches of intact ‘primary’ forest and is absent from secondary forest. Plants occur as scattered individuals at low density; they appear to be dioecious, female flowers are larger and fewer than male flowers. Pollinators are unknown. Fruits are 1–2 cm diameter, four-lobed orange berries, probably primate-dispersed (Cheek 2017).

###### CONSERVATION STATUS

*Vepris laurifolia* was only known from 18 individuals in Sierra Leone, but five of these are known to have been destroyed in recent years (hydroelectric dam flooding) with two probably destroyed (due to agriculture in the area) and six more are due to be lost in the next 1–10 years due to infrastructure developments (hydroelectric dam, transport corridor). Although only 11–13 surviving individuals are documented, it is possible that as many as 50–100 individuals may be found elsewhere, but intact forest habitat for this species only occurs as scattered remnants and is threatened with clearance for agriculture. Even in these scattered islands, the species is mostly absent (M. Cheek pers. obs. 2012–2016, Guinea; X. van der Burgt pers. obs. 2009–2016, Sierra Leone). It is also absent, or extremely rare, from most of Liberia where most of the surviving forest in West Africa remains. Botanical inventory work there over many years by C. Jongkind has not discovered this species (C. Jongkind pers. comm. to M. Cheek 2014). None of the large national parks (e.g. Tai National Park, Gola Rainforest National Park) are known to support it, despite botanical inventory effort. The species was assessed as Critically Endangered (CR) under criterion C2a(i) since less than 250 mature individuals are thought to exist and there is a continuing decline in the number of mature individuals, with less than 50 individuals in each subpopulation (Cheek 2017). In Guinea the species is included in two Tropical Important Plant Areas, Kounounkan and Ziama (Couch *et al*. 2019).

###### PHENOLOGY

Flowers mainly in April & May. Fruit in Sept.

###### ETYMOLOGY

Named for the resemblance of the leaves to those of the genus *Laurus* (Lauraceae)

###### VERNACULAR NAMES

Foh-foh-tae (fide Mamadou Camara of Oure Kaba cited in *Cheek et al*.16600). No uses are recorded.

###### NOTES

*Vepris felicis* was named by Breteler (1995) based on the Liberian specimen (*J.F.F. E. de Wilde* 3754, type specimen), the specimens collected in Guinea in 1937 near Mt. Benna (Jacques-Felix 2096 for from whom the species is named) and another specimen collected in 1954 near Mt. Kakoulima (*Schnell* 7568). Since the species was not named until 1995, it did not feature in the Rutaceae of Flora of West Tropical Africa (Keay 1958) and so is not mentioned in either Flore de la Republique de Guinée (Lisowski 2009) nor Flore de la Cote-D’Ivoire (Ake Assi 2001), both of which are based on the Flora of West Tropical Africa. Lachenaud and Onana (2021) recently discovered that *Garcinia laurifolia* was an earlier and unexpected synonym of *V. felicis*. The name *G. laurifolia*, was originally published in Clusiaceae, probably due to the opposite leaves and poor state of the type collection (*Scott-Elliott* 4806 from Ninia, Talla Hills, Guinea). Lachenaud and Onana (2021) have compared the types and there is no doubt that *G. laurifolia* is identical to *V. felicis* and since the epithet laurifolia is earlier and still available in *Vepris* it takes precedence, hence their publication of the new combination. The specimens we refer to above are additional to those given in Lachenaud & Onana (2021).

*Vepris laurifolia* is unusual among unifoliolate *Vepris* in that at the apex of flowering stems, the leaves are opposite (not alternate) and in that the leaflet is not articulated with the petiole, further the fruits are 4-locular (in other unifoliolate species they are 1 or 2-locular), with four widely separated style bases. This shrub is so unusual in its genus that flowering specimens in the field have been misidentified as *Rinorea* (Violaceae). However, the translucid spots usual in Rutaceae can be found using a lens, young leaves and bright light (Cheek 2017). This species should be a priority for molecular phylogenetic analysis since it is so morphologically anomalous.

##### 2. Vepris udzungwa

*Cheek* **sp. nov.** Type: Tanzania, Udzungwa Mountains National Park, Camp 357 – pt 358, 07.4° S, 36.37° E, 1980 m, fr. 12 Oct. 2002, *Luke et al*. 9109 (holotype K barcode K000875153! isotypes EA! NHT!, MO!). (Fig. 2)

**Fig. 2.**
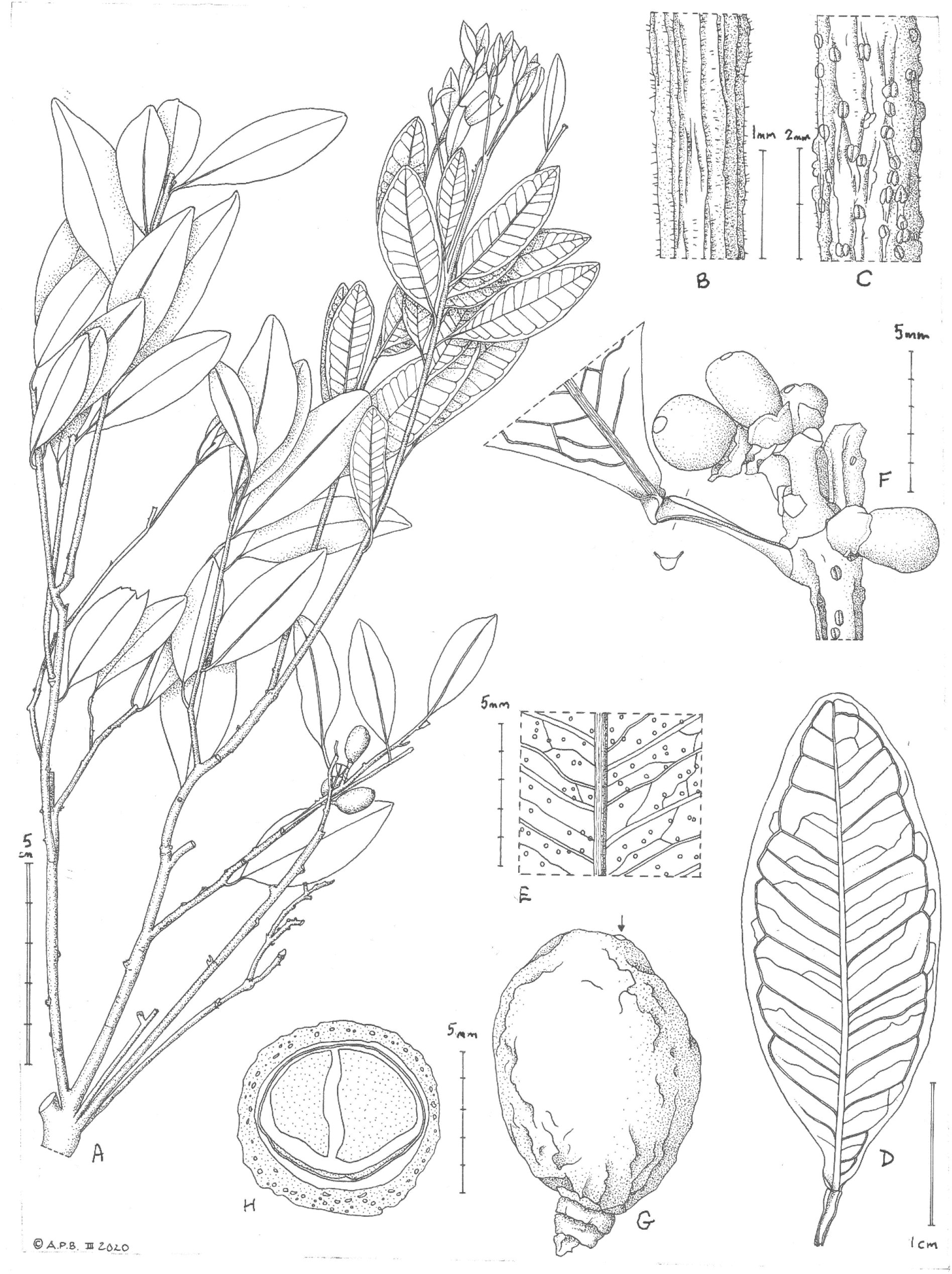
Vepris udzungwa. **A.** habit, fruiting branch; **B.** young stem showing hairs; **C.** older stem, showing dense lenticels and persistent hairs; **D.** leaf, adaxial surface; **E.** abaxial leaf surface showing conspicuous black oil gland dots; **F.** stem node and immature infructescence with leaf, showing winged petiole; **G.** fruit, side view, showing basal and apical asymmetry (stigma arrowed); **H.** transverse section of fruit showing seed. Scale bars: **A** = 5 cm; **D** = 1 cm; **E-H** = 5 mm; **C** = 2 mm; **B** = 1 mm. **A-E, G**&**H** from *Luke et al*. 9109; **F** from *Luke et al*. 8639. Drawn by Andrew Brown.

*Evergreen tree* 5 – 8 m tall, lacking scent when dried, densely branched. Leafy stem internodes (0.6 –)0.8 – 1.4(– 3.3) cm long, 1 – 2 mm diam., at the most distal leafy node, minutely puberulent when young, hairs white, simple, patent, c. 0.05 mm long, covering c. 10% of surface, glabrescent, epidermis rapidly becoming white-grey, densely (c. 50% of surface) lenticellate; lenticels raised, white, longitudinally elliptic, 0.75 × 0.5 mm, with a longitudinal midline groove. *Leaves* coriaceous, ± concolorous, dark green when live (*Luke et al*. 6895, K) drying green-yellow below, green-brown above, upper surface glossy, lanceolate-elliptic, less usually narrowly elliptic 3.7 – 6.8(– 7.3) x 1.6 – 2.5 cm, apex rounded, base broadly, convexly acute to subrounded, margin slightly revolute when dried; secondary nerves 7 – 8(– 10) on each side of the midrib, arising at c. 60° from the midrib, intersecondary nerves conspicuous, raised, forming a reticulum, tertiary and quaternary nerves not raised, less conspicuous; oil glands inconspicuous on upper surface, black and conspicuous on lower surface, (0 –)2 – 3(– 4) per mm^2^. *Petiole* articulated at apex, plano-convex, 0.3 – 0.6(– 0.9) cm long, 1.5 mm wide, margins with minute patent wings c. 0.4 mm wide, widest at articulation with blade, generally narrowing towards base, glabrous, crater-like glands inconspicuous. *Inflorescences* known from fruiting and post-anthetic material only, female inflorescences axillary racemes, 1 – 3(– 5)-flowered, inflorescence axis 3 – 10 mm long, glabrous. *Bracts* at base of pedicel, isodiametric, c. 4 × 4 mm, glabrous. *Pedicels* 1.5 mm long, glabrous.*_Sepals* 4, broadly triangular, c. 0.5 × 0.75 mm, glabrous. *Petals* not seen. *Stamens* (female flowers) 4, c. 1 mm long, filaments dorsiventrally flattened, tapering from base to apex, anthers orbicular, c. 0.3 mm diam.

*Fruit* ellipsoid to obovoid, slightly laterally compressed, 10 – 11 × 8 × 7 mm, asymmetric at base and apex, both pedicel and style inserted sublaterally on opposing sides, apex rounded, base truncate; style elliptic, flat, 1 mm wide; surface with irregular, longitudinal ribs, glabrous, surface oil glands inconspicuous; fruit wall 0.75 mm thick, endocarp not detected; uniloculate, 1-seeded. *Seed* ellipsoid 9 – 9.5 × 6.5 × 5.5 mm, seed coat dark brown, thinly leathery; embryo white, cotyledons equal, surface oil gland pits colourless.

###### RECOGNITION

*Vepris udzungwa* differs from *V. lukei* Cheek in the lateral nerves of the leaf-blade 7 – 8(– 10) on each side of the midrib (not 22 – 28), leaf apex rounded (not acuminate), and from all other E. African (Uganda, Kenya, Tanzania) unifoliolate species except *V. mildbraediana* in the hairy stems (not glabrous) and in the fruit asymmetric at both apex and base.

###### DISTRIBUTION

Tanzania, endemic to the summits of the Udzungwa Mts.

###### SPECIMENS EXAMINED

**TANZANIA. Udzungwa Mountains National Park**, Camp 357 – pt 358, 07.41° S, 36.37° E, 1980 m, fr. 12 Oct. 2002, *Luke et al*. 9109 (holo. K barcode K000875153!; iso: EA! NHT!, MO!); ibid, Luhomero Mt, camp 132 – 134, 07.47° S, 36.33° E, 2100 m, st. 3 Oct. 2000, *Luke et al*. 6895 (EA!, K, 000875155!); ibid, above exit gully, 07.40° S, 36.36° E, 2100 m, imm. fr. 1 June 2002, *Luke et al* 8639 (EA! K barcode K000875154!, NHT!, MO!).

###### HABITAT

Interface of montane evergreen forest with wet montane “grassland”; 1980 – 2100 m. alt. with *Asplenium rutifolium* (Bergius) Kunze (Aspleniaceae), *Vincetoxicum coriaceum* (Schltr.) Meve & Liede (Apocynaceae), *Psychotria cryptogrammata* E.M.A. Petit (Rubiaceae), *Diodella sarmentosa* (Sw.) Bacigalupo & E.L. Cabral (Rubiaceae), *Pauridiantha hirsuta* Ntore (Rubiaceae), *Geranium arabicum* Forssk. (Geraniaceae), *Crotalaria lukwangulensis* Harms (Leguminosae), *Coleus schliebenii* (Mildbr.) A.J. Paton (Labiatae), *Cyphostemma masukuense* (Baker) Wild & R.B. Drumm. ssp. *ferrugineo-velutinum* Verdc. (Vitaceae), *Cucumis oreosyce* H. Schaef. (Cucurbitaceae), *Peperomia retusa* (L.f.) A. Dietr. (Piperaceae), *Ranunculus multifidus* Forssk.(Ranunculaceae), *Eriocaulon transvaalicum N*.E. Br. ssp. *tofieldifolium* (Shinz) S.M. Phillips (Eriocaulaceae), *Satyrium crassicaule* Rendle (Orchidaceae), *Clutia abyssinica* Jaub. & Spach (Peraceae), *Cyanotis barbata* D. Don (Commelinaceae), *Fuirena stricta* Steud. ssp. *chlorocarpa* (Ridl.) Lye (Cyperaceae).

###### CONSERVATION STATUS

*Vepris udzungwa* is known from two specimen-sites at a single threat-based location, the well-protected (Q. Luke pers. obs.) Udzungwa Mountains National Park. The area of occupation is estimated as 8 km^2^ using the preferred IUCN cell-size of 4 km^2^. Extent of occurrence using Geocat is 17 km^2^.Therefore, we assess the species as Vulnerable VU D2, since although there are no immediate threats, should parts of the protected area be de-gazetted as has been proposed or has happened for other such areas in Tanzania (Qin *et al*. 2019) the risk is that the species would soon become lost to habitat clearance.

###### PHENOLOGY

Immature fruits in June, mature fruits and growth pulse (flush) in Oct.

###### ETYMOLOGY

Taking the name of the mountain range and National Park in which the species was discovered and to which it appears to be unique.

###### VERNACULAR NAMES & USES

None are known.

###### NOTES

*Vepris udzungwa*, apart from *V.mildbraediana*, is unique among all east African (Uganda, Kenya, Tanzania) unifoliolate *Vepris* species in the hairy stems (Fig. 1B), and also in the fruits which are not only asymmetric at the base (as in e.g. *V. hanaganensis*) but also at the apex, the style being subapical (Fig. 1G). In addition, alone among these species it has longitudinally irregularly ribbed wrinkled fruits (Fig. 1G).

*Luke et al*. 6895 had previously been determined as “?” and numbers 8639 and 9109 as “Vepris sp. *(=Luke* 6895)” by Kaj Vollesen indicating that in annotating the three specimens *Vepris sp. cf. eugeniifolia*, he recognised them to represent a possible distinct, unplaced taxon.

*Vepris udzungwa*, since it occurs at 2000 m. alt, has oblong-elliptic leaves 3.7 – 6.8 × 1.6 – 2.5 cm, which lack an acumen, and in which the fruits are subglobose, is most likely to be confused with the widespread *Vepris eugeniifolia* (Tanzania – Somalia) and *V. simplicifolia* (N. Malawi – Ethiopia). It differs from both in the puberulent stems (versus glabrous) and in the both basally and apically asymmetric fruit (versus symmetric), short (0.3 – 0.6(– 0.9) cm long), winged petioles (versus longer, canaliculate or cylindrical petioles).

An unusual feature of *V. udzungwa* is the very densely lenticellate stems. Parts of the older stems can be more than 50% covered in lenticels, while in most other *Vepris* species the older stems are only sparsely lenticellate. It is possible that *V. udzungwa* shares a recent common ancestor with *V. lukei* which occurs at the same mountain range, at a lower altitudinal band. However, these two species are morphologically unlikely to be confused (see under the last species, below).

Numerous other species have been relatively recently discovered and are restricted or are largely restricted to the Udzungwa Mts. e.g., *Polyceratocarpus askhambryan-iringae* A.J. Marshall & D. M. Johnson (Annonaceae, Marshall *et al*. 2016), *Trichila lovettii* Cheek (Meliaceae, Cheek 1989). *Ancistrocladus tanzaniensis* Cheek & Frim.-Møll. (Ancistrocladaceae, Cheek *et al*. 2000; Cheek 2000), *Lukea triciae* Cheek & Gosline (Annonaceae, Cheek *et al*. 2022b); *Uvariopsis lovettiana* Couvreur & Q. Luke (Annonaceae, Couvreur & Q. Luke 2010); *Toussaintia patriciae* Q. Luke & Deroin (Annonaceae, Deroin & Q. Luke 2005); *Vernonia luhomeroensis* Q. Luke & Beentje (Asteraceae, Luke & Beentje 2003); *Lijndenia udzungwarum* R.D. Stone & Q. Luke (Melastomataceae, Stone & Luke 2015)

##### 3. Vepris drummondii

Mendonça (1961: 84; 1963: 204); Type: Zimbabwe “S. Rhodesia, Melsetter Distr., Glencoe Forest Reserve, slopes of Mt Pene”, fl. 24 Nov. 1955, *Drummond* 4995 (K holotype barcode K000199467!, isotypes PRE barcode PRE0688690!, SRGH barcode SRGH0000250-0!)

###### DISTRIBUTION

This species is restricted to the southern foothills of the Chimanimani Mountains of Zimbabwe and Mozambique, and nearby Mt Pene and Tarka Forest Lands in Zimbabwe. Its presence in Mozambique was only confirmed in 2015, although there were earlier potential records (Timberlake *et al*. 2016). A record from Mt Mulanje in Malawi (specimen at Harare Herbarium) is considered to be erroneous and is omitted (Darbyshire *et al*. 2017)

###### REPRESENTATIVE SPECIMENS EXAMINED

Zimbabwe, Melsetter Distr., On Chambuka River bank above the hydroram, Tarka Forest Reserve, fl. Nov. 1970, *Goldsmith* 35/70 (K!, SRGH, WAG); ibid. Haroni River, confluence of Haroni and Timbiri Rivers., fr. April 1969, *Goldsmith*38/69 (BR, K!, SRGH, WAG).

###### HABITAT

This small puberulous shrub c. 0.7 m tall, is found in the deep shade of mixed evergreen forest, sometimes associated with rivers and gulleys, at low to mid-altitudes, 300 – 1,600 m.

###### CONSERVATION STATUS

*Vepris drummondii* is known from only 11 collections, collected between 1955 and 2015 and is listed as Vulnerable under criteria B1ab(iii)+2ab(iii) with an EOO of 69 km^2^ and an AOO of 32 km^2^ based on known occurrence data (Darbyshire *et al*. 2017). Although this may be a slight under-estimate, total AOO is unlikely to exceed 100 km2. Several of the localities for this species are within Forest Reserves, for example Glencoe Forest Land in Zimbabwe, although these are managed for commercial forest production rather than for biodiversity and so do not guarantee protection. At Maronga in Mozambique much of the lowland forest has been cleared outside of the Trans-Frontier Conservation Area core zone and there is also significant artisanal gold mining activity along the Mussapo River west of Maronga, which has almost certainly resulted in riverine forest loss there. However, there is still intact forest suitable for this species within the core protected area at Maronga (Darbyshire *et al*. 2017). The species is considered secure at both the Chimanimani National Park (Timberlake *et al*. 2016) and at Glencoe in Zimbabwe.

###### PHENOLOGY

Flowering in November, fruiting in April.

###### ETYMOLOGY

Named for the late Robert (Bob) Drummond (1924 – 2008), a life-long Africa botanist and botanical collector who collected the type specimen of this species and who was curator and a stalwart of the SRGH herbarium in Harare until the end of his life (Timberlake *et al*.2017).

###### VERNACULAR NAMES & USES

None are recorded.

###### NOTES

*Vepris drummondii* is unlikely to be confused with any other species since it is the only unifoliolate *Vepris* in the Flora Zambesiaca area (Mozambique, Malawi, Zambia, Zimbabwe, Botswana, Caprivi strip of Namibia). The other species are all trifoliolate. It is similar to *V. mildbraediana* of the Uluguru Mts of Tanzania, but that species has longer and denser hairs on the axes, and the partial-peduncles are only c. 2 mm long and few-flowered, while in *V. drummondii* they are much more fully developed. Nonetheless these two may be sister species to each other. “The shiny thin skinned deep red fruits resemble small cherries. Two to three seeds each fruit, green” (*Goldsmith* 38/69)

##### 4. Vepris mildbraediana

G.M. Schulze (in Mildbraed 1934:192; Kokwaro 1982: 23). Type: Tanzania, “Bezirk Morogoro, Uluguru Gebirge, Nordwestseite, Nebelwald”, fl. 8 Nov. 1932, *Schleiben* 2933 (Holotype B destroyed; isotype BR barcode BR000000627300!).

###### DISTRIBUTION

Tanzania, Uluguru Mts

###### REPRESENTATIVE SPECIMENS EXAMINED

Only known from the type specimen.

###### HABITAT

Submontane forest; 1860 m alt.

###### CONSERVATION STATUS

*Vepris mildbraediana* does not appear on iucnredlist.org. Since the collector of the type specimen stated that it was “isolated” we can deduce that only a single plant was seen. Given threats to habitats in the Uluguru Mts (Ndang’ang’a *et al*. 2007) and the record of only a single individual (and certainly less than 50) despite multiple surveys for plants (but not targeting this species so far as we know) we provisionally assess this species as CR B1(ab(i-iii)+B2ab(i-iii), D.

###### PHENOLOGY

Only known in flower in November. Fruits unknown.

###### ETYMOLOGY

Named for Mildbraed (Gottfried Wilhelm Johannes Mildbraed (1879 – 1954)), a heroic botanist who despite being captured in then German Kamerun in the first world war, losing all his specimens collected, as spoils of war to the British (they were sent to K), and being imprisoned in France 1914 – 1919, continued collecting specimens in tropical Africa (1907 – 1928) and as a taxonomist identifying and publishing his discoveries and those of others. He collected in Cameroon, Tanzania, Burundi and Rwanda among other places.

###### VERNACULAR NAMES

None are known.

###### NOTES

Kokwaro (1982) treated *Vepris mildbraediana* as an “Insufficiently known species”, stating that he had not seen the type nor any other specimens so named, and that *Bruce* 510, which he described as *Vepris sp. A*,“has many similar characters (and is from the same locality) except for its paniculate inflorescence. On the other hand, *V. mildbraediana* may be a synonym of *V. ngamensis* if its inflorescence is a raceme as stated.” Happily, thanks to JStor Global Plants and the African Plants Initiative (reference?), while the holotype at B is destroyed, an isotype at BR was detected and is available as a high quality image.

https://plants.jstor.org/stable/10.5555/al.ap.specimen.br0000006273002?searchUri=genus%3DVepris%26species%3Dmildbraediana

It shows that the type specimen is densely covered in long, patent, yellow-brown, hairs, persistent on the stem for 5 – 6 nodes, also on the petioles, abaxial midrib and inflorescence axis. No other unifoliolate East African species described has such dense indumentum. *Vepris udzungwa* is the only other African unifoliolate *Vepris* described that has hairy stems but those are only present at the first internode, and the hairs are white, appressed, sparse (c. 10% coverage of the surface) and minute (0.05 mm long). *Vepris mildbraediana* has a panicle, but the partial-peduncles are only c. 2 mm long, unlike the raceme reported for *V. ngamensis. Vepris sp. A* of FTEA, apart from being glabrous has more gracile inflorescence axes and the stamens are twice as long as the petals (in *Vepris mildbraediana* the axes are stout, and the petals as long as the anthers). There is no doubt that *Vepris mildbraediana* is a distinct species

##### 5. Vepris lukei

*Cheek **sp. nov.*** Type: Tanzania, Udzungwa Mountains National Park, 07.40°S, 36.39°E, Camp 366-pt 367 1800 m alt., fr. 15 Oct. 2002, *Luke W.R.Q. & P.A, et al*. 9166 (holotype K barcode K000875455!; isotypes: EA!, NHT!, MO!). (Fig. 3)

**Fig. 3.**
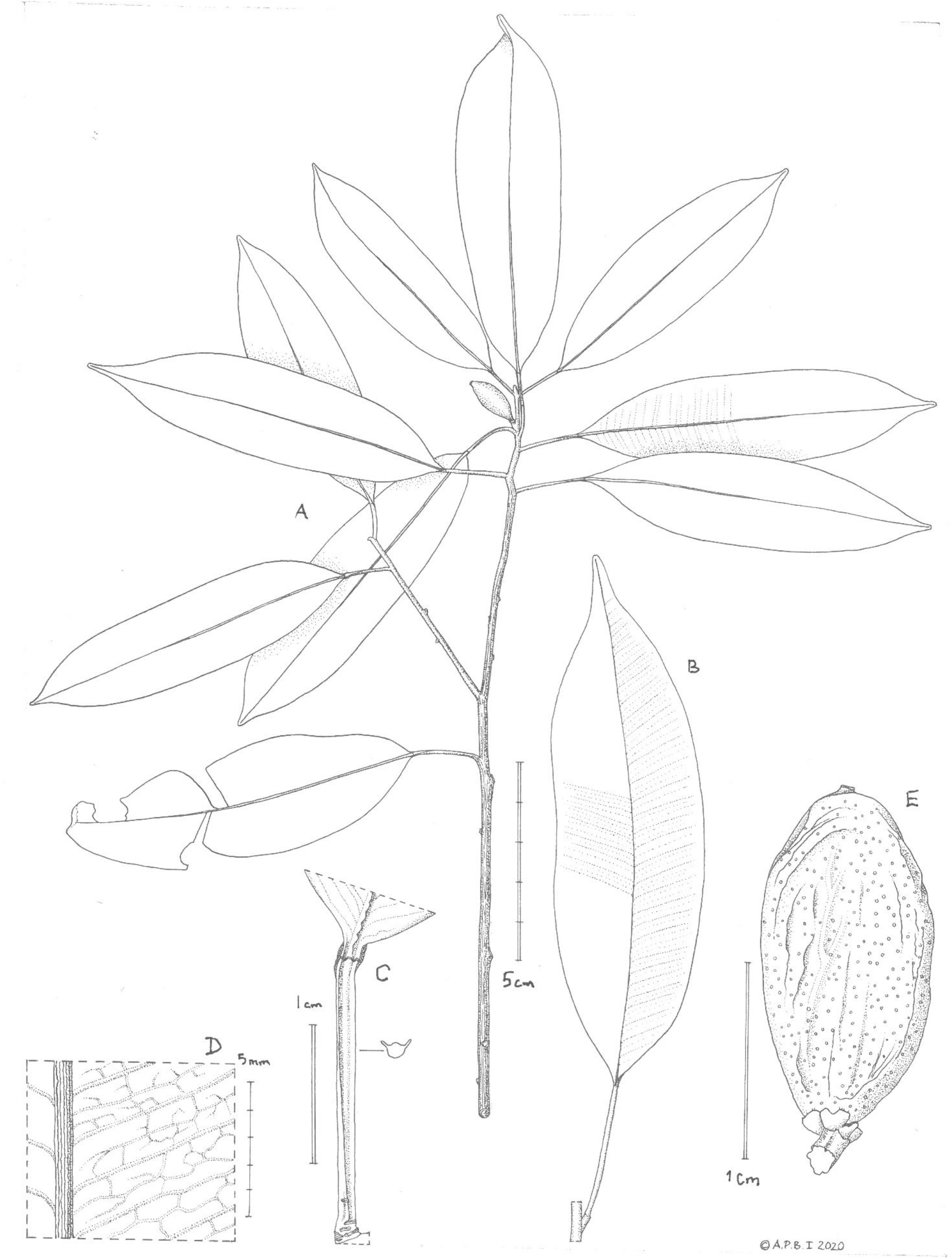
Vepris lukei. **A.** habit, fruiting branch; **B.** large leaf and attachment to stem; **C.** base of leaf-blade, articulation and winged petiole showing gland, together with transverse section of petiole to show wings; **D.** abaxial surface of leaf-blade showing reticulate quaternary nerves and inconspicuous oil glands; **E.** mature fruit showing raised black oil glands on surface. Scale-bars: graduated double bar = 5 cm; double bar = 1 cm; graduated single bar = 5 mm. **A** & **D** from *Luke et al*. 9166; **B, C,** & **E** = *Luke & Luke et al*. 10343. Drawn by ANDREW BROWN.

*Evergreen tree* 2 – 5 m tall, when dried smelling of dried fish. Leafy stems drying black, glossy, terete, 2 – 5 mm diam., internodes 4 – 26 mm long, increasing in length from the beginning of the flush (growth pulse), the main axis with 7 – 11 nodes per flush, growth of different flushes separated by 1 – 7 cm of naked stem; lenticels lacking, with fine longitudinal lines, glabrous. *Leaves* thinly coriaceous, ± concolorous, drying grey-green, glossy, narrowly oblong-elliptic, 6.5 – 13.7 × 1.8 – 3.7 cm, acumen 0.4 – 1.1 cm long, base broadly acute, margin undulate, slightly revolute, secondary nerves 22 – 28 on each side of the midrib, arising at c. 80° from the midrib, brochidodromous, forming a looping inframarginal nerve c. 2 mm from the margin; intersecondary nerves well developed, tertiary and quaternary nerves raised, forming a conspicuous reticulum on the lower surface; gland dots sparse, and barely detectable in transmitted light, only slightly translucent; concolorous so inconspicuous in reflected light except as minute raised spots on the abaxial surface, glabrous. *Petiole* articulated at apex, 2.5 – 48 mm long, variable in length; those first produced in a season longest, becoming successively shorter in successive nodes, plano-convex, c. 1 × 1 mm in section, the adaxial surface flat, the margins with a slender wing 0.5 mm wide, held at c. 45° from the vertical plane of the petiole axis, and bearing orbicular crater-like glands 0.2 – 0.25 mm diam., 2 – 8 mm apart. *Inflorescences* known from fruiting material only, female inflorescences 1 – 2 per stem, single, axillary in the leaf axils of the current season’s growth. *Bracts* 2, basal, opposite, connate, each forming a cupular pseudo-calyx 1 mm diam., c. 0.5 mm deep, glabrous. Pedicel 1(– 2) x 0.75 mm, glabrous. *Sepals* 4, triangular, c. 1 x 1 mm, becoming indurated in fruit, glabrous. *Fruit* ripening orange, cylindric-ellipsoid, 1-seeded, 15 – 17 x 7 – 10 mm, stigma remains subglobose, apex flattened, 0.5 mm long, 0.75 mm diam., surface with raised black oil glands 0.2 mm diam., 2 – 3 per mm^2^, glabrous. Pericarp 0.5 mm thick, endocarp vascularised, adhering to epicarp. *Seed* the same shape and slightly smaller than fruit, testa pellicular, brown; cotyledons 2, equal, the outer surface white, pitted with oil glands 0.1 mm diam.

###### RECOGNITION

*Vepris lukei* Cheek is similar to *Vepris robertsoniae*, differing in the 1-fruited infructescence, fruit surface with conspicuous, large black, raised oil glands, and fruit apex rounded (vs 2 – 5-fruited, surface with inconspicuous minute or absent oil glands, fruit apex acute to slightly rostrate), petioles winged, 0.25 – 4.8 cm long (vs canaliculate, (0.35 –)0.5 – 1.8(– 2.8) cm long), secondary nerves 22 – 28 each side of the midrib, stem epidermis black (vs secondary nerves 8 – 15 each side of the midrib, stem epidermis becoming dull white). Additional diagnostic characters can be found in table 1.

**Table 1.** Diagnostic characters separating *Vepris robertsoniae* from *Vepris lukei*

|  | *Vepris robertsoniae* | *Vepris lukei* |
| --- | --- | --- |
| Stem epidermis (dried material) | Pale brown, ageing dull white | Black, persisting black with age |
| Number of secondary nerves on each side of the midrib | 8 – 15 | 22 – 28 |
| Visibility of oil gland dots on abaxial leaf-blade surface | Conspicuous, black | Inconspicuous, concolorous with blade. |
| Petiole length | (0.35 – )0.5 – 1.8( – 2.8) cm | 0.25 – 4.8 cm |
| Petiole shape | Canaliculate, wings absent | Winged |
| Fruit apex | Acute to slightly rostrate | Rounded |
| Fruit surface: oil glands (dried material) | Inconspicuous, minute, concolorous or absent | Conspicuous, large black, raised. |
| Infructescence | 2 – 5-fruited | 1-fruited |
| Habitat | Lowland semi-evergreen forest, usually on limestone; 0 – 290m alt. | Submontane evergreen forest on crystalline rocks; 1540 – 1800m alt. |
| Geography | S.E. Kenya (Lamu, Kwale, Kilifi Districts) | Udzungwa Mts, Tanzania |

###### DISTRIBUTION

Tanzania, Udzungwa Mts.

###### SPECIMENS EXAMINED. TANZANIA

**Udzungwa Mountains National Park**, 07.40°S, 36.39°E, Camp 366-pt 367 1800 m alt., fr. 15 Oct. 2002, *Luke, W.R.Q. & P.A et al*. 9166 (K holo!; iso EA!, MO!, NHT!); Ndundulu Forest Reserve, 07.47°S, 36.39°E, Camp 589 1540m alt., fr. 6 Sept. 2004, *Luke et al*. 10343 (EA!, K!, MO!).

###### HABITAT

Submontane forest; 1540 – 1800 m alt. Associated species (identifications of *Luke et al*. specimens collected with *Vepris lukei*): *Hypoestes forskaolii* (Vahl) R. Br. ssp. *forskaolii, Sclerochiton uluguruensis* Vollesen (Acanthaceae), *Isolona linearis* Couvreur, *Monodora glob flora* Couvreur (Annonaceae), *Vincetoxicum anomalum* (N.E. Br.) Meve & Liede (Apocynaceae), *Diospyros* sp. *Luke & Luke* 9165, 9166 (Ebenaceae), *Erythrococca sanjensis* Radcl.-Sm. (Euphorbiaceae), *Streptocarpus kirkii* Hook.f. (Gesneriaceae), *Jasminum abyssinicum* DC. (Oleaceae), *Ixora scheffleri* K. Schum. & K. Krause ssp. *scheffleri, Pauridiantha paucinervis*(Hiern) Bremek. *ssp. holstii* (K. Schum.) Verdc. *Psychotria cryptogrammata* E.M.A. Petit, *Tarenna roseicosta* Bridson, *Tricalysia aciculiflora* Robbr. (Rubiaceae), *Vepris stolzii* I. Verd., *Zanthoxylum gilletii* (De Wild.) P.G. Waterman (Rutaceae), *Dracaena fragrans* (L.) Ker Gawl. (Dracaenaceae), *Aframomum laxiflorum* Lock (Zingiberaceae).

###### CONSERVATION STATUS

*Vepris lukei* is known only from the two specimens cited above, both shown on Google earth as being within the Udzungwa Mts National Park (although the southern site, formerly in the Ndundulu Forest Reserve is now officially the West Kilombero Nature Reserve contiguous with the National Park, Q. Luke pers. obs.), separated by c. 13 km. The protection level of this National Park is high currently, and so threats do not exist for this species at present (Q. Luke pers. obs. 2002 – 2004). However, with only a single location, a low number of individuals (only two specimens were seen despite hundreds of specimens being collected over two years by Luke and associates, and before and since by other botanical field workers) and an area of occupation estimated as 8 km^2^ using the preferred IUCN cell-size of 4 km^2^ (extent of occurrence cannot be calculated with only two points), this species would have a significant risk of extinction (from clearance of the forest habitat for agriculture as has been widespread in E Arc Forests) were the current excellent management levels to be lowered or the area to be partly degazetted as a National Park as has been proposed for other protected areas in Tanzania e.g. Serengeti National Park and Selous Forest Reserve (Qin *et al*. 2019). Since the species fits the IUCN (IUCN Standards and Petitions Committee 2022) definition of VU D2 “of a species not declining, but ..characterized by an acute restriction in their area of occupancy or in their number of locations thereby rendering them particularly susceptible to a plausible threat” we assess *Vepris lukei* as VU D2 (Vulnerable).

###### PHENOLOGY

Leaf flushing in September. Fruiting in September and October, flowering unknown.

###### ETYMOLOGY

Named for William Richard Quentin Luke, better known as Quentin Luke (1952-), lead collector of all known specimens of *Vepris lukei,_*and the most prolific living field botanist in East Africa. He is a Kenyan botanist and is Research Associate of the East African herbarium (EA). Full biographical and bibliographical information can be found in Polhill & Polhill (2015: 276 – 277). He has brought to light previously unknown species from across Africa e.g., in eastern Democratic Republic of Congo: *Keetia namoyae* O. Lachenaud & Q. Luke (Lachenaud *et al*. 2017) and from Mali and Guinea the only endemic African *Calophyllum, C. africanum* Cheek & Q. Luke (Cheek & Luke 2016; Couch *et al*. 2019). He has discovered numerous new species of plants especially in Kenya and Tanzania, such as the incredible spectacular Tanzanian tree acanth *Barleria mirabilis* I.Darbysh. & Q.Luke (Darbyshire & Luke 2016). He has also collected and described many other novel plant species from Tanzania and Kenya. More than ten species are named for him, e.g. *Keetia lukei* Bridson (Rubiaceae, Bridson 1994), including also the Tanzanian species *Cola quentinii* Cheek (Cheek & Dorr 2007) and *Cola lukei* Cheek (Cheek 2002). Most recently *Lukea* Gosline & Cheek, a new genus to science has been named in his honour (Cheek *et al*. 2022b).

###### VERNACULAR NAMES

None are known.

###### NOTES

*Luke et al*. 10343 had previously been identified as *Vepris robertsoniae* ined., and *Luke & Luke* 9166 as “*Vepris* sp., not matched” by Kaj Vollesen in 2004.

*Luke et al*. 10343 has new shoots with expanding leaves, and also the leaves from the previous season’s growth. These show a progressive reduction in length of the petiole during a season’s growth. The first formed petiole is 48 mm long, the second formed 43 mm long, then 38 mm, 30 mm, 25 mm, 22 mm, 14 mm, 10 mm, 5 mm, and finally at the end of the growth pulse, before dormancy, 2.5 mm long.

*Vepris lukei* has many similarities with *V. robertsoniae* and for this reason they may share a recent common ancestor and may well be sister species. Both species smell of fish when dried, have numerous parallel secondary nerves, crater glands on the petiole, are glabrous except for the sepal margins, lack panicles and nectar discs. For these reasons it is logical that material of *V. lukei* was formerly named as *V. robertsoniae*. However apart from ecology and geography, the two species differ in several key morphological characters (Table 1) and there is no doubt that they are distinct.

*Vepris lukei* is unusual amongst E African unifoliolate *Vepris* species in possessing winged petioles. All other species have canaliculate or terete petioles. It also is unusual in the extreme high number of secondary nerves, 22 – 28 on each side of the midrib – resembling a *Calophyllum*. Further, it is unique in this group in the highly reduced female inflorescences which appear to be 1-flowered. Examination of immature fruiting specimens gives no indication that they bore more than one flower. Yet this species remains known from only two collections, and male and female flowers at anthesis remain to be obtained.

The geographical and ecological disjunction between the two very similar and probably sister species, one at low altitude in the coastal forests of SE Kenya, the other at high altitude in the Eastern Arc Mountains of Tanzania, is seen in several other genera, such as *Lukea*, with *L. quentinii* Gosline & Cheek in Kenyan coastal forest, and *L. triciae* in the Udzungwa Mts (Cheek *et al*. in press), *Ancistrocladus* Wall. with *A. tanzaniensis* Cheek & Frim.-Møll. in the Udzungwas and *A. robertsoniorum* J.Léonard in the Kenyan coastal forests (Cheek *et al*. 2000, Cheek 2000), also in the genus *Afrothismia* with *A. mhoroana* Cheek in the Ulugurus and *A. baerae* Cheek in Kenyan coastal forests (Cheek 2004b; Cheek 2006; Cheek & Jannerup 2006). Numerous other taxa are restricted to the Eastern Arc Mts of Tanzania and the Kenyan Coastal Forests, which together are referred to as EACF (see discussion).

New plant species are still steadily being discovered for science and published from Tanzania, other recent examples being *Mischogyne iddii* Gosline & A.R. Marshall (Annonaceae, Gosline et al. 2019), *Hibiscus hareyae* L.A.J.Thomson & Cheek (Malvaceae, Thomson & Cheek 2020), *Inversodicraea tanzaniensis* Cheek (Podostemaceae, Cheek *et al*. 2020a) and *Keetia davidii* (Rubiaceae, Cheek & Bridson 2019),

##### 6. Vepris robertsoniae

*Q. Luke & Cheek **sp. nov.*** Type: Kenya, Kwale District, Marenji, 50 m, fl.,18 Dec. 1990, *W.R.Q. Luke & S.A. Robertson* 2679 (holotype K barcode K000875137!; isotypes EA!, MO!, UPPS!). (Fig. 4 – 7).

**Fig. 4.**
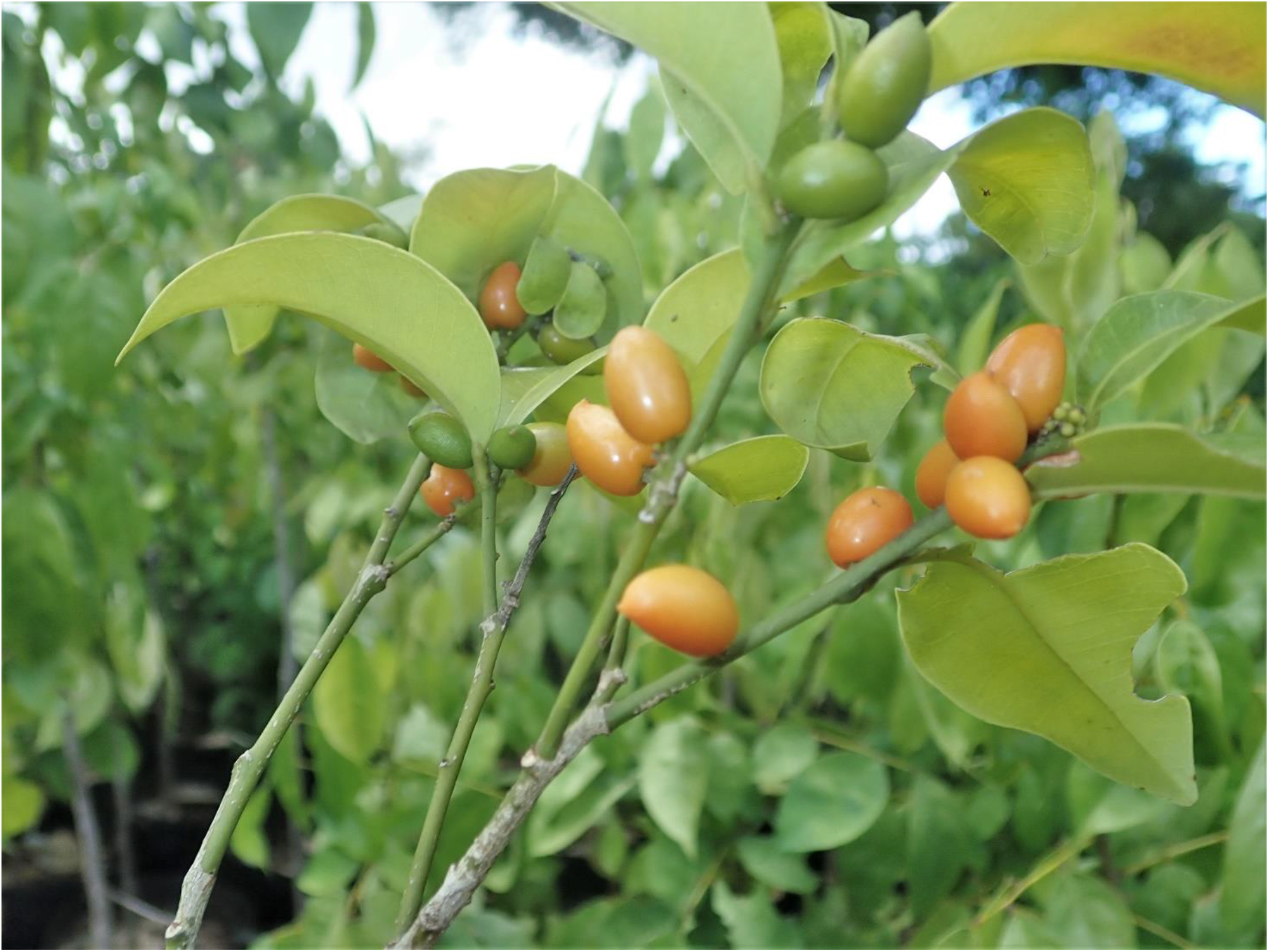
Vepris robertsoniae. Habit of fruiting shrub, Base Titanium nursery 15 April 2019. Photo by W.R.Q. Luke

**Fig. 5.**
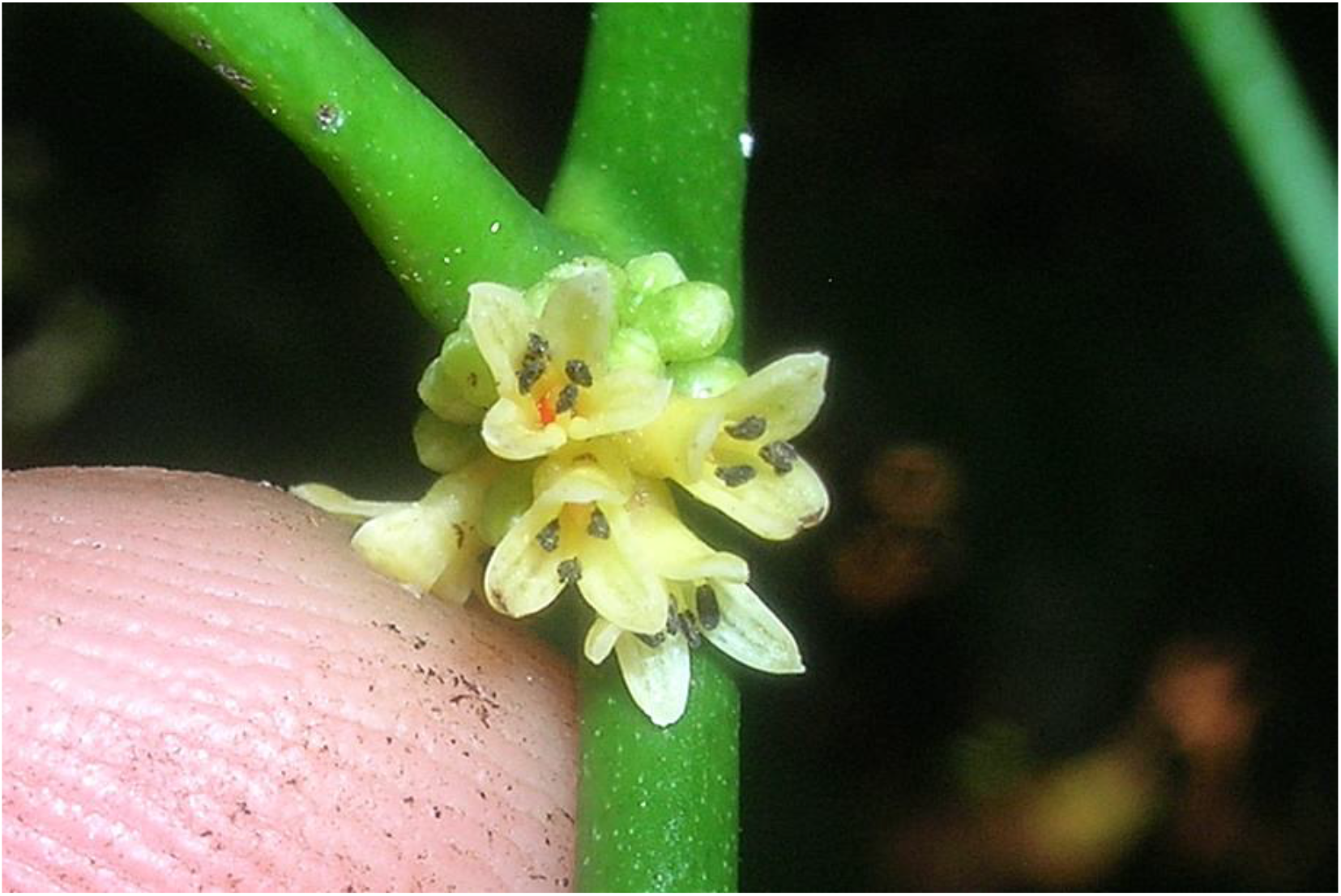
Vepris robertsoniae. Close up of male flowers, note the four stamens, cultivated plant in Base Titanium nursery 28 May 2021. Photo by W.R.Q. Luke

**Fig. 6.**
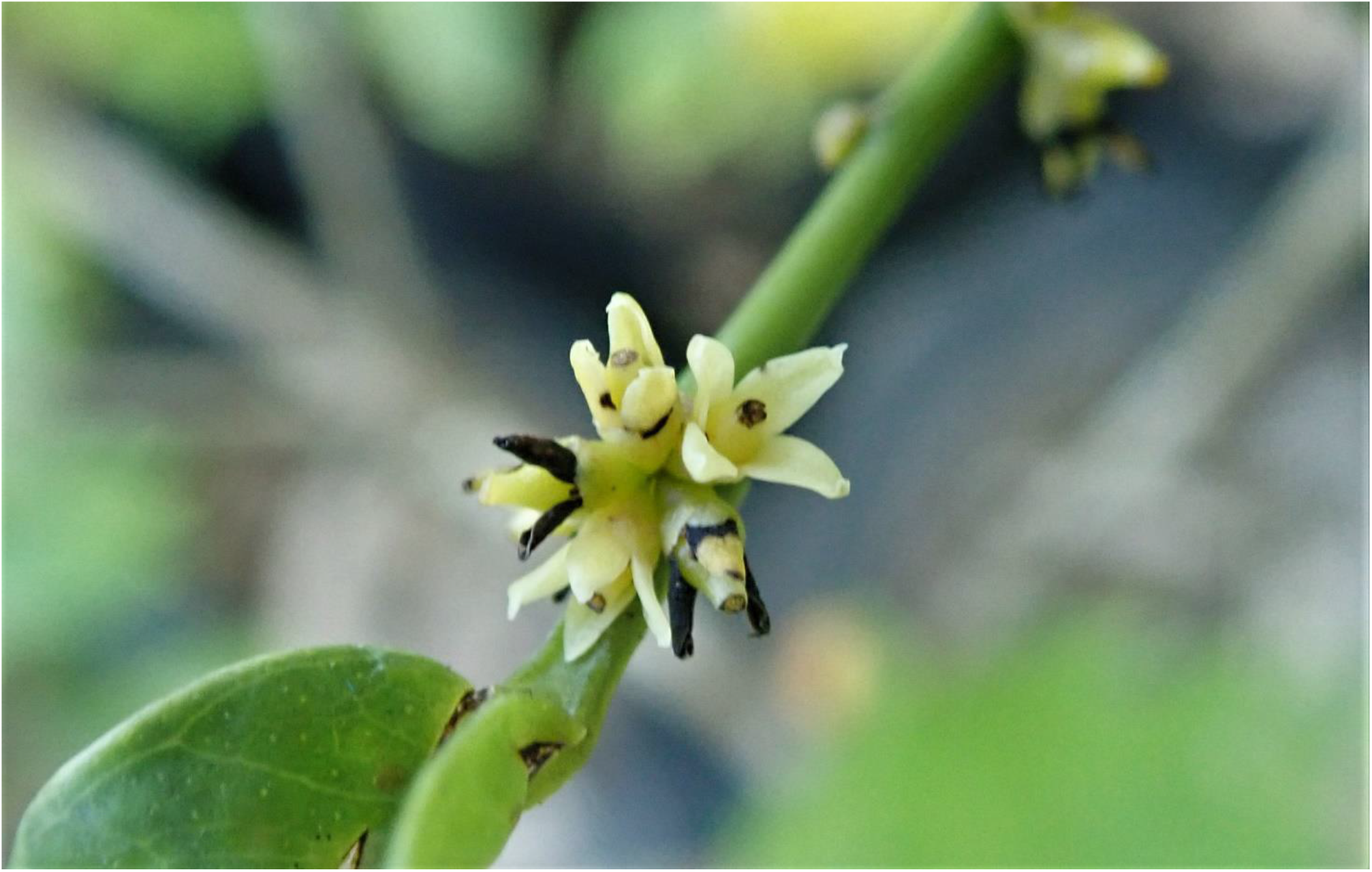
Vepris robertsoniae. Close up of female flowers, 11 Nov. 2020, Base Titanium nursery. The Eight staminodes are concealed. Photo by W.R.Q. Luke

**Fig. 7.**
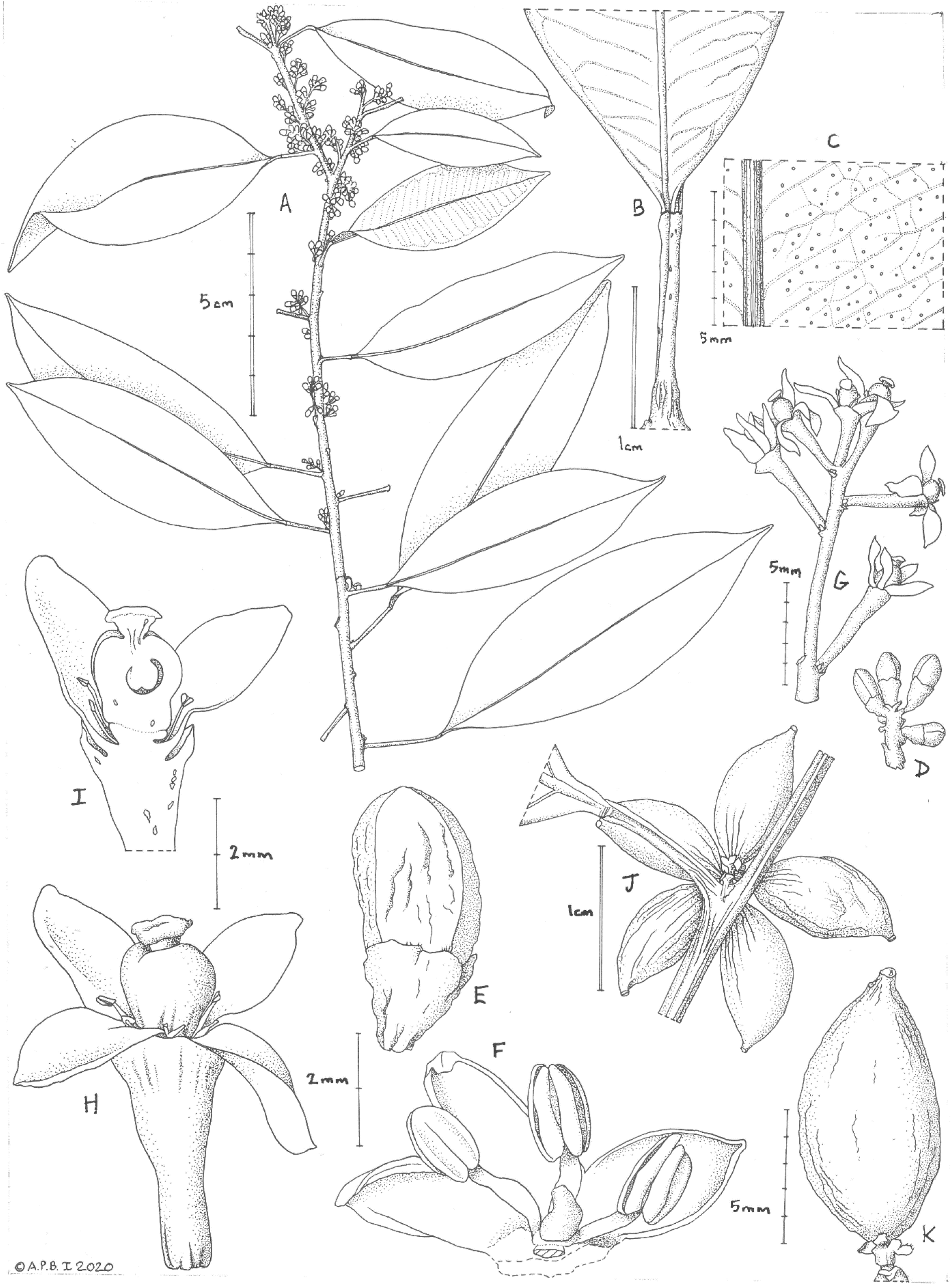
Vepris robertsoniae. **A.** habit, flowering branch; **B.** base of leaf-blade showing articulation with canaliculate petiole and glands; **C.** abaxial surface of leaf-blade showing surface with black oil glands; **D.** male inflorescence; **E.** male flower bud; **F.** male flower with stamen and petal removed to show pistillode; **G.** female inflorescence; **H.** female flower; **I.** longitudinal section of female flower showing unilocular ovary with a single pendulous ovule. **A, B, D-F** from *Luke et al*. 2679; **C** from *Luke et al*. 1670; **G-I** from *Robertson* 6852; **J, K** *Luke et al*. 1771; Drawn by ANDREW BROWN.

*Vepris robertsonae* Q.Luke *ined*. (Luke 2005: 62)

*Small evergreen tree or shrub* (1.5 –)2 – 6m tall, dried specimens smelling of dried fish. Leafy stems drying glossy brown-green, finely longitudinally wrinkled, terete, internodes (0 –)0.8 – 4(– 9.3) cm long, 2 – 4(– 5) mm diam. at the lowest leafy node, becoming pale, whitish grey, lenticels rare, sparse, white longitudinally elliptic, 0.3 – 1.4 × 0.2 – 0.4(– 0.5) mm *Leaves* thickly coriaceous, drying grey-green above, almost concolorous, but the lower surface slightly brown, elliptic, rarely slightly ovate-elliptic, (3.75 –)7 – 13.2(– 18.4) x (1.5 –)3 – 5.1(– 6.35) cm, acumen short and broad, (0 –)0.4 – 1.2(– 1.8) cm long, sometimes absent, base broadly acute or rounded; margin slightly revolute when dry; secondary nerves 8 – 15 on each side of the midrib, arising at 40 – 50° from the midrib, brochidodromous, forming a looping inframarginal nerve c. 2 mm from the margin; intersecondary nerves as well developed as secondary nerves, tertiary nerves reticulate, the nerves raised; gland dots clear and bright in transmitted light, about 1(– 3) per mm2, in reflected light conspicuous, black, but not raised on the abaxial surface, 0.2 mm diam., (0 –)1 – 2(– 4) per mm2; glabrous. *Petiole* articulated at apex, longest produced at start, shortest at end of growth season, canaliculate, (0.35 –)0.5 – 1.8(– 2.8) cm long, 1 – 1.5 mm wide, the ventral groove slit-like, margins with scattered crater-like glands. *Inflorescences* 8 – 15 per leafy stem, 4 – 10-flowered, racemose, axillary, 0.5 – 1.4 cm long, peduncle 1 – 1.5 mm long, bracts 0.1 mm long. *Male flowers* with pedicel c. 0.5 mm long. Sepals 4, quadrangular, 0.3 – 0.6 × 0.8 – 1 mm, glabrous apart from a few simple marginal hairs 0.05 mm long. Petals 4, elliptic-oblong, 3.5 × 1.5 – 1.75 mm, apex slightly acuminate. Stamens 4, c. 3 mm long, filaments 1.5 mm long; anthers ellipsoid 1.5 – 1.75x 1 mm, glabrous. Disc absent. Pistillode 1 x 0.6 mm, glabrous. *Female flowers* with pedicels (2.5 –)3 – 4 mm long, dilated at apex, sepals 4 as in male flowers. Petals oblong-elliptic, 2.5 – 2.8 × 1.8 mm, apex obtuse. Staminodes 8, c. 1 mm long, 4 shorter than others, with vestigial anthers. Ovary obovoid, c. 2 mm long, proximal third 1 mm diam., unilocular, distal two-thirds 1.5 mm diam., apex retuse, style c. 0.3 mm long, widening from base to apex, stigma peltate, c. 1 mm diam. *Infructescence* 2 – 5-fruited. *Fruits* yellow-orange (live), 1-seeded, ellipsoid or ovoid-ellipsoid 8 – 11.5 × 4 – 5.5(– 7.5) mm, apex weakly rostrate or acute, rostrum c. 1 mm long, base rounded, pericarp leathery, thin, surface lacking oil glands, glabrous. *Seed* ellipsoid-ovoid c. 9 – 5 mm, encased in endocarp. Endocarp cartilaginous, translucent, laced with a network of flattened vascular bundles, brown; seed-coat membranous; cotyledons equal, white, surface black, pitted with oil glands c. 0.1 mm diam.

###### RECOGNITION

Similar to *Vepris eugeniifolia* (Engl.) I. Verd., differing in the elliptic (rarely slightly ovate-elliptic) leaf-blades (vs ovate); flowers single along the rhachis in the inflorescences (vs glomerules along the rhachis); fruits ovoid-ellipsoid or ellipsoid, apex acute or slightly rostrate (vs globose, apex rounded). Additional diagnostic characters are given below in the notes and in table 2.

**Table 2.** Diagnostic characters separating *Vepris robertsoniae* from *Vepris eugeniifolia*.Characters for *Vepris eugeniifolia* taken from Kokwaro (1982).

|  | *Vepris eugeniifolia* | *Vepris robertsoniae* |
| --- | --- | --- |
| Scent of dried leaves | Odourless | Dried fish |
| Leaf-blade shape | Ovate or lanceolate | Elliptic (rarely slightly ovate-elliptic) |
| Leaf-blade dimensions | 3.5 – 9 x 2 – 4.2 cm | (3.75 – )7 – 13.2( – 18.4) x (1.5 – )3 – 5.1( – 6.35) cm |
| Inflorescence length | 1 – 3( – 8.5) cm | 0.5 – 1.4 cm |
| Number of styles (both female and male flowers) and locules | 2 | 1 |
| Fruit shape | Globose, apex rounded | Ellipsoid or ovoid-ellipsoid, apex slightly rostrate or acute |
| Petiole | Cylindric (terete) | Canaliculate |
| No. secondary nerves each side of the midrib | 6 – 8( – 9) | 8 – 15 |
| No. stamens in male flowers | 8 | 4 |

###### DISTRIBUTION

Coastal Kenya: Lamu, Kilifi and KwaleCounties.

###### SPECIMENS EXAMINED. KENYA

**Lamu District,** Lunghi Forest Reserve, 35 m, st., 1 Dec. 1988, *W.R.Q. Luke & S.A. Robertson* 1539 (EA!, K000875135!); **Kilifi District,** between Dzitsoni & Jaribuni, 150 m, fl., 21 Feb. 1989, *WR.Q. Luke & S.A. Robertson* 1670 (EA!, K000875136!); Kilifi District, Mangea Hill, 450 m, fl., 25 March 1989, *W.R.Q. Luke & S.A. Robertson* 1824 (EA!, K000875143!); Kilifi District, Mangea Hill (Sita), 290 m, fr., 24 Mar 1989, *W.R.Q. Luke & S.A. Robertson* 1771 (EA!, K000875144!, MO!); Kilifi District, Kaya Jibana, SW slope, 200 m, fl., 14 Dec. 1990, *WR.Q. Luke & S.A. Robertson* 2642 (EA!, two sheets: K000875145!, K000875146!); **Kwale District,** Marenji, 50 m, fl.,18 Dec. 1990, *WR.Q. Luke & S.A. Robertson* 2679 (holo. K000875137!; iso. EA!, MO!, UPPS!); Kwale District, Mwachi Forest Reserve NW corner and down to Mwachi river, 0.359° S 39.32° E, 30-80 m, fl.,17 May 1990, *S.A. Robertson & W.R.Q. Luke* 6187 (EA!, two sheets: K000875138!, K000875139!, MO!); Kwale District, Mwaluganji Forest Reserve (including Kaya Mtae), 0.405° S 39.27° E, 200 – 300 m, fl., 15 Nov. 1989, *S.A. Robertson & W.R.Q. Luke* 6044 (EA!, K000875140!) Kwale District, Gongoni Forest, 30m, st., 3 June 1990, *W.R.Q. Luke & S.A. Robertson* 2395 (EA!, K000875141!); Kwale District, Gongoni Forest, 30 m, fl., 9 June 1990, *WR.Q. Luke* 2415 (EA!, K000875142!); Kwale District, Gongoni Forest, 30 m, fl., 9 June1990, *W.R.Q. Luke* 2416 (EA!, two sheets: K000875147!, K000875148!, UPS!); Kwale District, Diani forest, 0.420° S 39.34° E, 5 m, fl., 29 Aug. 1993, *S.A. Robertson* 6852 (EA!, K000875149!).

###### HABITAT

Lowland semi-evergreen forest, usually (always?) on limestone; 5 – 200 (– 290) m alt. Associated species (identifications of *Luke et al*.specimens collected with *Vepris robsertsoniae): Ecbolium amplexicaule* S. Moore, *Thunbergia stelligera* Lindau, *Trichaulax mwasumbii* Vollesen (Acanthaceae), *Psilotrichum majus* Peter (Amaranthaceae), *Solanecio angulatus* (Vahl) C. Jeffrey (Compositae), *Dictyophleba lucida* (K. Schum.) Pierre (Apocynaceae), *Diospyros shimbaensis* F. White (Ebenaceae), *Triaspis mozambica* A. Juss. (Malpighiaceae), *Eugenia verdcourtii* Byng (Myrtaceae), *Afrocanthium kilifiense* (Bridson) Lantz, *Cladoceras subcapitatum* (K. Schum. & K. Krause) Bremek., *Coffea pseudozanguebariae* Bridson, *Coptosperma supra-axillare* (Hemsl.)

Degreef, *Didymosalpinx norae* (Swynnerton) Keay, *Pavetta crebrifolia* Hiern var. crebrifolia, *Psydrax faulknerae* Bridson, *Rothmannia manganjae* (Hiern) Keay, *Rytigynia parvifolia* Verdc., *Tricalysia pallens* Hiern (Rubiaceae), *Haplocoelum inoploeum* Radlk. (Sapindaceae), *Rinorea squamosa* (Tul.) Baill. ssp. *kaessneri* (Engl.) Grey-Wilson (Violaceae), *Cyphostemma zimmermannii* Verdc. (Vitaceae), *Zamioculcas zamiifolia* (Lodd.) Engl.(Araceae), *Angraecum teres* Summerh., *Calyptrochilum christyanum* (Rchb.f.) Summerh. (Orchidaceae)

###### CONSERVATION STATUS

*Vepris robertsoniae* is known from nine locations with an extent of occurrence of 7825 km^2^ and area of occupation of 88 km^2^. Threats include degradation of habitat, such as cutting of poles. Therefore the species was assessed as Vulnerable, VU B1ab(i,ii,iii,iv,v)+2ab(i,ii,iii,iv,v) (Musili *et al*. 2020).

###### PHENOLOGY

Flowering Nov.-June (-Aug.), fruiting in March-June.

###### ETYMOLOGY

Named for Mrs Anne Robertson of Kenya, pioneering collector of plants and early champion of the conservation of Kenya’s coastal forests where her studies resulted in the discovery of plant species several others of which, apart from *Vepris robertsoniae* are named for her, including*Barleria robertsoniae* I.Darbysh. (Darbyshire *et al*. 2010) and *Psydrax robertsoniae* Bridson (Bridson 1991). Anne Robertson also produced a checklist of the Seychelles Islands, collecting there also, and is commemorated there by. *Cynanchum robertsoniae* Liede (Liede 1995). Finally, she is commemorated with her late husband Ian by the coastal Kenyan forest liana species *Ancistrocladus robertsoniorum* J. Léonard (Ancistrocladaceae, Léonard 1984; Cheek 2000).

###### VERNACULAR NAMES

None recorded.

###### NOTES

*Vepris robertsoniae* is most likely to be confused with *Vepris eugeniifolia* (see table 2) which also occurs at low altitudes on coral rock in S.E. Kenya, is probably sympatric, but which is much more common and widespread (Tanzania to Somalia).

Apart from both species being unifoliolate and glabrous, with similar gland dots they also share the key character formerly ascribed in FTEA to distinguish *Vepris* in the former narrow sense: 8 stamens. But this number of stamens are only present in the female flowers of *V. robertsoniae:* the male flowers have four stamens. Characters separating the two species are given in Table 2. The two are easily separated vegetatively since *V. robertsoniae* has a canaliculate (not terete) petiole, the blade has 8 – 15 lateral nerves (not 6 – 8(– 9)) on each side of the midrib. The base of the blade is broadly acute or rounded, the margin at that point convex or straight, while in *V. eugeniifolia* it is usually concave. In dried specimens of the last species the blade folds along the midrib, exposing the abaxial surface in old leaves on a sheet, while those of *V. robertsoniae* remain flat.

*Vepris robertsoniae* has also been confused with *V. lukei*. See under the last species for a discussion of their affinities and for diagnostic characters separating them (Table 1).

Numerous other species with a similar range to *Vepris robersoniae*, also threatened and restricted to a set of Kenyan coastal Kaya forests, have been steadily documented in recent decades. Examples include *Uvariodendron dzomboense* Dagallier, W.R.Q. Luke & Couvreur (Kaya Dzombo, EN B1ab(iii)+2ab(iii)) and *Uvariodendron schmidtii* W.R.Q. Luke, Dagallier & Couvreur (Longomwagandi, likely VU)(both Annonaceae, Dagallier *et al*. 2021), *Croton kinondoensis* G.W.Hu, V.M.Ngumbau & Q.F.Wang (Kaya Kinondo, likely CR, Euphorbiaceae, Ngumbau *et al*. 2020), *Saintpaulia ionantha* H.Wendl. ssp. *rupicola* (B.L. Burtt) I. Darbysh. (Cha Simba, CR A2ac, B2ab(i,ii,iii,iv,v), Gesneriaceae, Darbyshire 2006), *Keetia lukei* Bridson (Kaya Jibana and Gongoni F.R, EN B1ab(iii)+2ab(iii), Rubiaceae, Bridson 1994) and *Premna mwadimei* Ngumbau & G.W. Hu (Cha Simba, CR B1ab(iii)+2ab(iii), Labiatae, Ngumbau *et al*. 2021).

Cultivated plants, collected as seedlings from Gongoni Forest Reserve in July 2014, began to flower and fruit when they attained about 1.2 m tall after four to five years (observations from the Base Titanium nursery in coastal Kenya by the second author). The planting medium used was a sandy soil mix with coir and manure. Flowering occurs April-June, Nov. & Dec, fruiting April -June, and December.

##### 7. Vepris welwitschii

(Hiern) Exell (1929: 148; Exell & Mendonça 1951: 272; Figueiredo & Smith 2008: 155). Type: Angola, “in montibus petrosis supra Tandambando”, fr. Nov. 1854, *Welwitsch*471 (Lectotype LISU, barcode LISU206243!, syntypes BM barcode BM00798355!, LISU barcode LISU206244!, PRE barcode PRE0601859-0!“Zenzo do Golungo”)

*Glycosmis welwitschii* Hiern (1896: 115)

*Vepris gossweileri* I. Verd. (1926: 399) non Mziray (1992: 72). Angola. Type: Serra do Socollo-Undui, between Ambriz and Lifuni River, **“** Loanda, Cazengo”, fr. 11 Dec. 1907, *Gossweiler* 4895 (Holotype K, barcode K000199522!; isotypes COI barcode COI COI00040965!, K barcode K000199523!).

###### DISTRIBUTION

Angola. The species is only known from a few specimens in Bengo and Cuanza Norte provinces in northwestern Angola. It is known from four localities (Lachenaud & Onana 2021).

###### REPRESENTATIVE SPECIMENS EXAMINED

Angola, Luanda: Icala e Bengo – Macchias de Catete, fr. 1929, *Gossweiler* 9173 (COI barcode COI00040964!)

###### HABITAT

*Vepris welwitschii* is restricted to xerophytic vegetation on limestone outcrops up to 800 m in altitude (Lachenaud & Onana 2021).

###### CONSERVATION STATUS

*Vepris welwitschii* was assessed as Near Threatened by Timberlake (2021b), stating that it is not widely distributed and that only historic records are available since it was last recorded in 1921. Timberlake stated that it has an extent of occurrence (EOO) of 8,368 km^2^ and an area of occupancy (AOO) of 20 km^2^ calculated from the four known collecting localities and that there appears to have been land cover change from agriculture and settlement at some of the localities that could threaten the species. In contrast to Timberlake, Lachenaud & Onana (2021) assess the species as Endangered EN B2ab(iii) citing an EOO of 14,092 km^2^, and AOO of 12 km^2^ and an expected decline due to habitat clearance for e.g. charcoal. The second assessment appears to better reflect the extinction risk status of the species.

###### PHENOLOGY

Flower buds and immature fruits in September, mature fruits in October (Lachenaud & Onana 2021).

###### ETYMOLOGY

Named for the Austrian, Friedrich Welwitsch (1806 – 1872), the most famous botanical collector of specimens in Angola, who collected the original specimens from which the species was described. He is also commemorated by the genus *Welwitschia* Hook.f. (Hooker 1863).

###### VERNACULAR NAMES & USES

None are known

###### NOTES

*Vepris welwitschii* a tree to 6 m tall, is most likely to be confused, and indeed has been, with *V. africana*, the only other unifoliolate species of the genus that occurs in Angola. The two can be distinguished using the characters cited above under the second species and in the key to species. Most notably *Vepris welwitschii* has black fruit, not orange or red as is usual in the genus. Lectotypification, synonymy and delimitation of this species was expertly performed by Lachenaud & Onana (2021). However, they opted to choose as lectotype a syntype at LISU for which there is no evidence that Bentham, credited author of the name, had seen. The syntype at BM does not have this deficiency. They also point out that this species remains incompletely known e.g. open flowers are not available.

##### 8. E eugeniifolia

(Engl.) I. Verd. (Verdoorn 1926: 399; Kokwaro 1982: 17; Beentje 1994: 371; Thulin 1999:177; Friis 1992: 184, fig. 115; Luke 2005: 62). Type: Tanzania, Usambara Mts, Mashewa (« Mascheua »), 500 m, fl. Aug. 1893, *Holst* 8869 (B holotype probably destroyed; isotypes BM, G barcode G00445210!, HBG barcode HBG510346!, K barcode K000199492!, M barcode M-0110250!, S sheet number 08-9780!)

*Toddalia simplicifolia* var. *eugeniifolia* Engl. (Engler 1895: 228) *? Teclea gracilipes* Engler (1917: 308). Type: Tanzania, Uzaramo Distrct, *Stuhlmann* 1894 (B holotype probably destroyed)

*Aegelopsis alexandrae* Chiov. (Chiovenda 1932: 50). Type: Somalia, Giubia, isola Touata di Alexandra, July 1931, *Zozzi* 327, (isotype K barcode K000199447!)

*Teclea alexandrae* (Chiov.) Senni (1935: 82)

###### DISTRIBUTION

Ethiopia, Somalia, Kenya, Tanzania

###### REPRESENTATIVE SPECIMENS EXAMINED

**ETHIOPIA.** 12 km NE of Telte towards Brindi and Yavello, 1150 m alt., fr. 24 Nov. 2010, *Friis et al*. 13882 (ETH, K!); **SOMALIA.** 20 km from Fanoole barrage. Jess site 54. st. 31 Jan. 1988, *Deshmukh* in Jess 435 (K!); Summit of Bur Juqalalan, 300 – 630 m, fr. 30 Feb. 1982, *Beckett* 1700 (K!). **KENYA.** Northern Prov., Dandu, fr. 11 April 1952, 800 m, *Gillett* 12761 (EA, K!); West of Malindi, N bank of Galana River, st. 13 Feb 1953, *Woodley in Bally* 8586 (K!); Makueni Distr., Kibwezi FR, 975 m alt., fr. *Luke* 14376, EA, K!), Kilifi, fl. 23 Dec. 1936, *Moggridge* 221 (EA, K!). **TANZANIA.** Genda-Genda South, fr. 27 June 1982, *Hawthorne* 949 (EA, FHO, K!); Handeni Distr., Kwa Mkono, 600 m, fr. 20 Feb. 1980, *Archbold* 2737 (DSM, EA, K!)

###### HABITAT

Coastal forest and semi-evergreen shrubland on coral rag or normal soil, or at higher altitudes in *Acacia-Commiphora* woodland, rainfall ranges 500/1000 mm p.a. (e.g. Friis 1992: 185); 0 – 630 (– 1827) m alt.

###### CONSERVATION STATUS

*Vepris eugeniifolia* does not appear oniucnredlist.org, but from its wide range and numerous sites it is likely to be assessed as Least Concern.

###### PHENOLOGY

Fruits June, Nov.-Dec. in Ethiopia, Feb. in Somalia, Apr.-Aug. in Kenya & Tanzania. Flowering Dec.-Feb. in Ethiopia, May(-July) in Somalia, Dec.-April, July-Oct. in Kenya and Tanzania.

###### VERNACULAR NAMES & USES

Agnio golet *(Zozzi* 327, K!), filfil owliyi *(Deshmukh* in Jess 435, K!), rehdo *(Beckett* 1700, K! all Somali, Somalia); Mwaowa (Wakulu) leaves boiled in water and administered orally for canine complaints (Kenya, Kilifi *Moggridge* 221, K!), root bark used in the preparation of arrow poison (W Malindi, *Woodley in Bally* 8586, K!)

###### NOTES

Not rarely confused with the usually higher altitude *V. simplex* especially at mid to low altitudes in Kenya and Ethiopia. While in Ethiopia *V. simplex* grows at altitudes of 1900 – 2000 m in *Podocarpus* forest, *Vepris eugeniifolia* grows in drier and lower habitats e.g. 1100 – 1400 m alt., in *Acacia-Commiphora* woodland, and in fact can survive in drier habitats than any other African unifoliolate *Vepris*, witness that it is the only unifoliolate species to occur in Somalia (Thulin 1999). The leaves are acuminate (usually rounded in *V. simplex’)* and their size range is smaller, although the largest leaves of *V. eugeniifolia* can exceed the smallest of *V. simplex*. The flowers are extremely different, those of *V. simplex* being twice the size and having four not eight stamens, the females with one style not two, and the fruits of *V. simplex* are smaller 3 – 4(–5) mm diam., subsessile, drying black or orange, while those of *V. eugeniifolia* are 6 – 8 mm diam., drying with a white waxy layer on 4 – 6 mm long pedicels.

##### 9. V. sp. A of FTEA

Kokwaro (1982:18); Mziray (1992: 78)

###### DISTRIBUTION

Tanzania, Morogoro Distr.

###### SPECIMENS EXAMINED

Tanzania, Morogoro Distr., Uluguru Mts, Bunduki, fl. 10 Jan 1935, *Bruce* 510

###### HABITAT

Submontane forest c. 1700 m alt.

###### CONSERVATION STATUS

*Vepris sp.A* of FTEA has not been formally named and therefore does not appear on iucnredlist.org. Provisionally it should be regarded as Critically Endangered CR D since only a single plant is known at a site that has threats (Ndang’ang’a *et al*. 2007). Forest loss at Uluguru Mts has been concentrated in the habitat of *Vepris sp.A* of FTEA (see discussion).

###### PHENOLOGY

Flowering in January, fruits unknown.

###### VERNACULAR NAMES

None are recorded

###### NOTES

Kokwaro (1982: 18) recognised this entity and stated “The specimen is somewhat similar to *Teclea amaniensis* except the stamens are clearly 8. It is however, inadequate to formally describe a new species. It is also close to *Vepris ngamensis* but here the inflorescence is a panicle. See also *V. mildbraediana*, p. 23”. Treated by Mziray (1992: 78) as an “Insufficiently known taxon”. This entity appears to be a most distinct and yet undescribed species.

##### 10. Vepris amaniensis

(Engl.) Mziray (pro parte 1992: 70)

Types: Tanzania, Amani, *Engler* 565 (Syntype, B destroyed)*; Warnecke* 516 (Syntype B, destroyed); Neotype proposed here Tanzania “Tanganyika Terr., Amani”, 5 April 1922, *Salmon* 171 K barcode K000593352!; EA)

*Teclea amaniensis* Engl. (Engler 1905: 244; Kokwaro 1982: 24 pro parte)

*Vepris ngamensis* Engl. ex Verdoorn (1926: 399); Kokwaro (1982: 18). Type: Tanzania, E. Usambara Mts, Amani, *Engler* 565 (holotype B destroyed; neotype selected here: Tanzania “Tanganyika terr., Amani, 4 April 1919, *Salamani bin Kilwa* G6172 (Neo. K barcode K000593351!; isoneo EA)). **synon. nov.**

###### DISTRIBUTION

Tanzania, Muheza Distr., Usambara Mts at Amani and Bulwa.

###### SPECIMENS EXAMINED

TANZANIA. Muheza Distr. Amani, *Engler* 565 (B syn. destroyed);ibid. *Warnecke* 516 (B syn., destroyed); ibid. Amani, 5 April 1922, *Salmon* 171 (K neo.!, EA isoneo.); ibid Amani, 4 April 1919, *Salamani bin Kilwa* G6172 (K neo.!, EA isoneo); ibid. Amani, Urwald, fr. 22 July 1911, *Grote* AH 3416 (K!); E. Usambara, Bulwa, Ukundo, imm. fr. 27 Aug. 1980, *Kibuwa* 5342 (K!); ibid. old fl., fr. 27 Aug. 1980, *Kibuwa* 5343 (K!); ibid, just below Amani, 2900’, fl. 20 March 1950, *Verdcourt* 122 (K!, two sheets); ibid., Amani Forest, near the guest house, fr. 3 Aug. 1986, *Lovett, Ellis, Keeley* 869 (K!, MO).

###### HABITAT

*Vepris amaniensis* is a 0.5 – 3 m tall shrub in evergreen forest with *Myrianthus, Allanblackia* (Clusiaceae), *Memecylon cogniauxii (*Melastomataceae, *Verdcourt* 122), *Cephalosphaera usambarensis* (Myristicaceae)*, Anisophyllea obtusifolia* (Anisophylleaceae, *Lovett et al*. 869); 900 – 1000 m alt.

###### CONSERVATION STATUS

Timberlake (2021a) in assessing the extinction risk of *V. amaniensis*states:.Some of the forests from which *Vepris amaniensis* is recorded, particularly in Tanzania, are under threat of clearance for small-scale and subsistence agriculture. The extent of occurrence (EOO) is calculated at 210,887 km^2^ and the minimum area of occupancy (AOO) is 104 km^2^. As there are only nine recorded locations the species is assessed as Vulnerable VU B2ab(ii,iii,v). However, it has recently been discovered (see Notes below) that this species is restricted to near Amani and Bulwa in the Usambara Mts, with a far smaller AOO and EOO, and so will merit reassessment, likely as EN.

###### PHENOLOGY

Flowering in March and April, fruits in July and August.

###### ETYMOLOGY

Meaning “from Amani”, referring to the origin of the original specimens which were collected at or near Amani in the Usambara Mts of then German E Africa, Tanganyika, now Tanzania.

###### VERNACULAR NAMES & USES

None are recorded.

###### NOTES

While finalising the key and skeletal species accounts for this paper, the first author found that the specimens assigned to this species at K, although concordant as a whole with the description in Kokwaro (1982), contained more than one species. Most of the material was not in agreement with the description in the original protologue of Engler (1905), nor the description by Verdoorn (1926), which appears based on Engler’s description (although is less precise). It seems that between the time of Verdoorn (1926), who only cited *Warnecke* 516K, and Kokwaro (1982), numerous additional specimens of at least one other unifoliolate shrub was collected in the Usambaras and adjoining areas, including Kenya, and erroneously attributed to *V. amaniensis*, although accommodated in the expanded description of the species in FTEA. Most of this material has the apex of the petiole winged, hairy stems, an inflorescence shorter than the petioles, and often the odd trifoliolate leaf among the predominantly unifoliate ones. These seem to represent a further new species that will be the subject of a future paper. None of the specimens of the putative new species were collected in Amani. In contrast, only seven surviving specimens (see specimens examined above) represent a species that fits the descriptions of Engler (1905) and of Verdoorn (1926). These have thin papery, elliptic leaflets with a length: breadth ratio of c. 2.5:1, glabrous stems, petioles which are terete at base and canaliculate at apex, inflorescences 0.9 – 4(– 5) cm long, far exceeding (usually) the petioles, and leaves which are uniformly unifoliolate. All the specimens are from Amani except two from nearby Bulwa. Since they match Engler’s protologue description and location, a neotype has been selected from among them that matches the original description, since all the original material of *V. amaniensis* (the syntypes *Engler* 565 and *Warnecke* 516k in Herb. Amani) have been destroyed or lost. Although Mziray (1992) states the last is at K there is an ancient annotation to a species cover that this specimen is “not here”. In addition, the label of Salmon G 6171 (E African Agricultural research station, Amani, 5 April 1922) states in script contemporary with the original label, “The type is not in herb. Amani”. This suggests that no duplicates were left by Engler’s team in the Amani Herbarium (so they could not have been transferred to EA with the rest of that herbarium).

*Vepris ngamensis* is here formally added to the synonymy of the earlier published *V. amaniensis*.Treated by Mziray (1992: 78) as an “Insufficiently known taxon”, *Vepris ngamensis* is only known from certainty from the type, *Engler* 565, also collected at Amani, but destroyed at Berlin. Although Kokwaro also attributed *Drummond & Hemsley* 3349 (not found, presumed missing) to *V. ngamensis* he had not actually seen the original material. When Verdoorn described *Vepris ngamensis* in 1926 from material that had been annotated by Engler as *Teclea ngamensis* (Verdoorn 1926; Kokwaro 1982) she presumably missed the fact that this same specimen is one of the two syntypes of *V. amaniensis*. Comparing the original descriptions of *Vepris ngamensis* (Verdoorn 1926) with that of *V. amaniensis* Engler (1905) shows no point of morphological difference except in the number of stamens. The first having four (hence assigned to the genus *Teclea)* and the second seven (so ascribed to *Vepris*). While some specimens cited above have four stamens, another *(Salamani bin Kilwa)* is annotated “Stamens 5 – 6!”. Although stamen number was used to assign species to different genera formerly, and has value as a species character, *Mziray* (1992) cited the range in variation of stamens from 4 to 8 (sometimes on the same plant) in *V. heterophylla* as evidence that this is not in itself a reliable character for generic separation, nor even in some cases for separating species. We neotypify *V. ngamensis* above, in the absence of any original material, choosing material from the type location that matches its protologue most closely.

##### 11. Vepris africana

(Hook.f. ex Benth.) O.Lachenaud & Onana (2021: 109). Type: S.Tomé, without date or locality, *Don s.n*. (Holotype K, barcode K000199556). (Fig. 8)

**Fig. 8.**
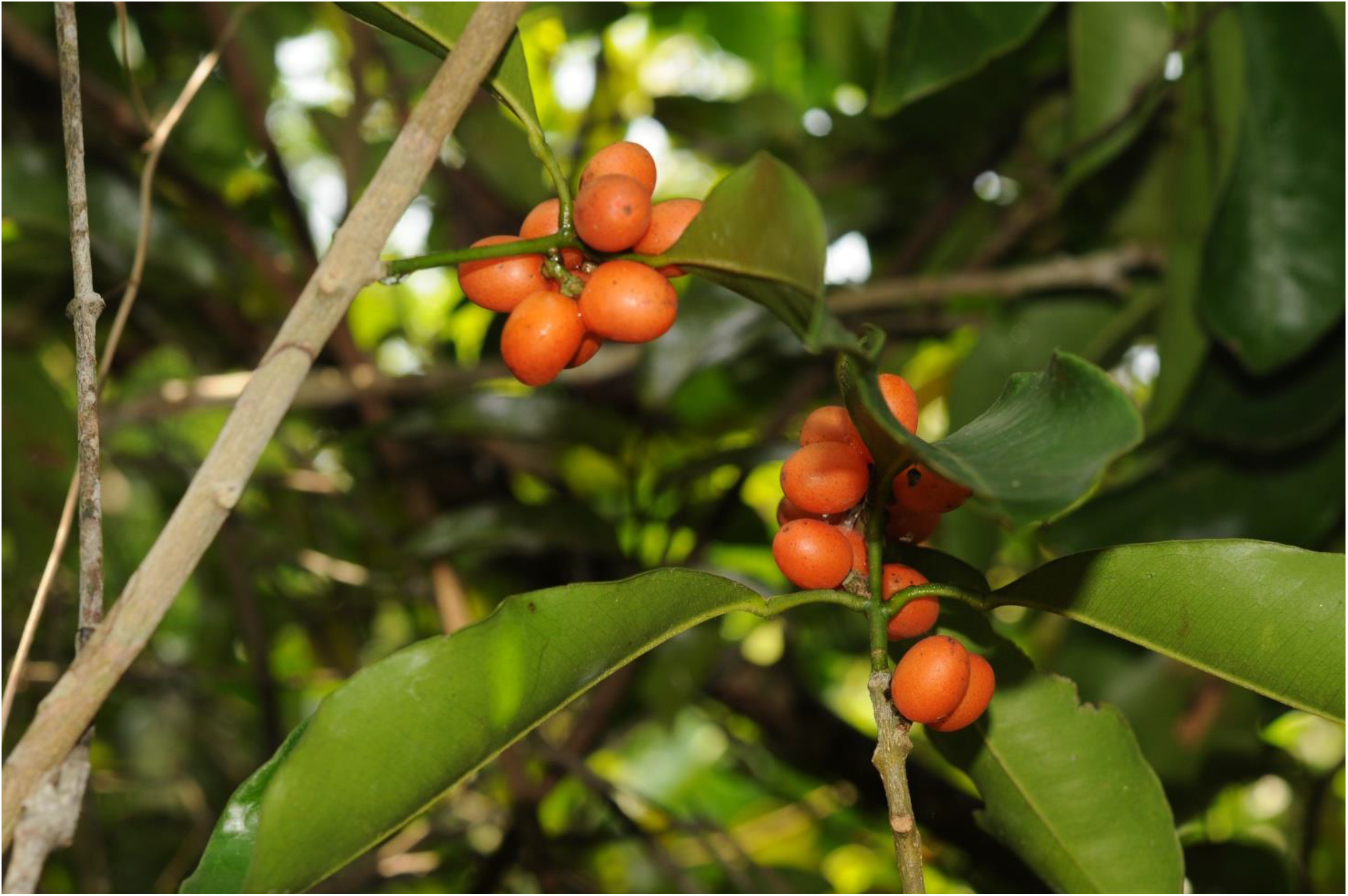
Vepris africana. Habitat of fruiting plant (*Mpandzou* 1282A, IEC, K) in white sand coastal thicket near Pointe Noire, Republic of Congo in 2012. Photo by M. Cheek.

*Glycosmis? africana* Hook.f. ex Benth. in W. J. Hooker (1849: 256).

*Teclea gossweileri* I. Verd. (1926: 409); Exell & Mendonça (1951: 271). Type: Angola, Cuanza Norte, Cabiri, 1 July 1921, *Gossweiler* 8328 (Holotype K, barcode K000199528, K000199529) *Vepris gossweileri* (I. Verd.)Mziray nom. illeg. Mziray (1992:72; Figueiredo & Smith 2008: 155; Langat *et al*. 2021)

###### DISTRIBUTION

N. Angola (both metropolitan and Cabinda), Republic of Congo, Gabon and S. Tomé

###### REPRESENTATIVE SPECIMENS EXAMINED

**Republic of Congo.** Bas - Kouilou, a 1.5 km du pont Bas - Kouilou, au bord de la RN5, fr. 13 Dec. 2012, *Mpandzou* 1906 (IEC, K); Pointe Noire, fr. 10 July 2011, *Mpandzou* 1282A (IEC, K); Tchimpounga Point 1 zone soleil 1, fr. 13 Nov. 2012, *T. Kami* 1327 (IEC, K, MO).

###### HABITAT

Coastal thicket, often on white sand (then sometimes with *Vepris teva* in Congo), forest patches in wooded grassland, sometimes in rocky areas, sometimes on limestone; sea-level-1200 m alt. (Lachenaud & Onana 2021 pro parte)

###### CONSERVATION STATUS

*Vepris africana* does not appear on iucnredlist.org. including under its synonyms. Lachenaud & Onana (2021) give a detailed provisional conservation assessment stating that it is Near Threatened based on 22 herbarium specimens, five of which could not be placed geographically, resulting in 11 IUCN threat-based locations, an AOO of 48 km^2^, an EOO of 369,480 km^2^. Threats observed by Lachenaud in Gabon and S. Tomé are habitat loss and degradation from charcoal production, urbanisation, and agriculture. The first author has observed these same threats, and additionally port construction facing the species in the Republic of Congo, where already locations have been lost, and others are set to follow. Given this data the authors opt to assess the species as Vulnerable VU D2 since they agree with Lachenaud & Onana (2021) that further of the known locations are likely to disappear in the near future.

###### PHENOLOGY

Flowers (June-) Sept.-Jan.; fruit Nov. and Feb.

###### ETYMOLOGY

Named for Africa by J. Hooker.

###### VERNACULAR NAMES & USES

None are known.

###### NOTES

Lachenaud & Onana (2021) resolved the nomenclature of this taxon and give an excellent and detailed description, ecological and other notes and original line drawing of this species which we have drawn upon here, supplemented by the first author of this paper’s original observations of the species in Republic of Congo and of live plants observed in cultivation. Plants grow relatively easily and vigorously from seed but even after 10 years had not flowered (Langat *et al*. 2021 under the synonym *Vepris gossweileri*). Laboratory investigation has shown multi-layered anti-bacterial synergism in combinations of minor compounds with E-caryophyllene in this species (Langat *et al*.2021). In the Republic of Congo the species is only known from a distinctive coastal thicket on white sand where it can grow with *Vepris teva* (Gosline *et al*. 2014; Cheek *et al*. 2014; Langat *et al*. in press). Lachenaud & Onana (2021) report that *Vepris africana* is unusual for the genus in being androdioecious, pollen is produced in both flower types but male flowers have pistillodes only. This feature separates it from the similar but dioecious Comorian *V. unifoliolata* (Baill.) Labat, M. Pignal & O. Pascal They attribute the presence of the species on S.Tome a volcanic oceanic island, as probably resulting from dispersal by frugivorous birds or possibly by marine currents. The specimens cited above are additional to those reported in Lachenaud & Onana (2021) but do not increase the range of the species.

*Vepris africana* has been confused with *V. welwitschii* in Angola where they both occur, and both species are superficially very similar. However, the first has subsessile flowers (pedicels 0 – 0.5 mm long), unilocular ovaries and 4 stamens, orange fruits, the second pedicels 1 – 2.5 mm long, is bilocular, has 8 stamens, and black fruits.

##### 12. V. *hanangensis*

var. ***unifoliolata*** Kokwaro (1978: 791). Type: Kenya, Nairobi, Karura Forest, fr. 23 Jan. 1970, *Perdue & Kibuwa* 10241 (holotype EA barcode EA000003105!; isotypes BR barcode BR0000006273699!, K barcode K000199486!, PRE barcode PRE0594695-0!)

###### DISTRIBUTION

Kenya, only known from Karura Forest of Nairobi

###### REPRESENTATIVE SPECIMENS EXAMINED

Kenya, outskirts of Nairobi, Karura Forest, 25 Oct. 1976, *Kokwaro* 4038 (EA, K).

###### HABITAT

Upland dry evergreen forest; c. 1700 m alt.

###### CONSERVATION STATUS

*Vepris hanangensis* var. *unifoliolata* is listed as Vulnerable (World Conservation Monitoring Centre 1998b) under criterion D2, listing urbanisation and land clearance for agriculture as among the key threats. In the last 20 years, Nairobi has expanded greatly, reducing and degrading habitat. However, due to a successful campaign led by Wangari Maathai to reject all allocations of land in Karura, and subsequent fencing by the local residents association, the habitat of this highly range-restricted taxon is protected and an assessment of Vulnerable VU D2 seems appropriate. It is advisable that there is a baseline survey to verify that the taxon survives, and against which monitoring and a management plan for the tree can be devised

###### PHENOLOGY

Fruits are known in January.

###### ETYMOLOGY

Named for the unifoliolate leaves of the mature trees that distinguish this taxon from the typical variety of the species which is trifoliolate.

###### VERNACULAR NAMES

None are known.

###### NOTES

Young plants of this variety frequently have some 3—foliolate, some 2-foliolate and a majority of 1-foliolate leaves. Unifoliolate leaves from young plants are exceptionally large, up to 30 x 12 cm (Kokwaro 1978: 791).

*Vepris hanangensis* var. *unifoliolata* in leaf might be confused with *V. simplex* which also occurs at this altitude. However, *Vepris hanangensis* var. *unifoliolata* as in the typical variety, has long cylindrical fruits held in large persistent panicles, unlike the globose fruits on reduced racemes of the other species.

The collectors of the type stated that the tree grew up to 150 feet (=45 m) tall. This would make it by the far the tallest growing of the African unifoliolate *Vepris* species. However, this is an error since the tallest tree in Karura Forest is no more than 15 m tall (QL pers. obs. 2022). Only the two specimens cited are known to us.

##### 13. Vepris simplex

Cheek. nom. nov. (Fig. 9).

**Fig. 9.**
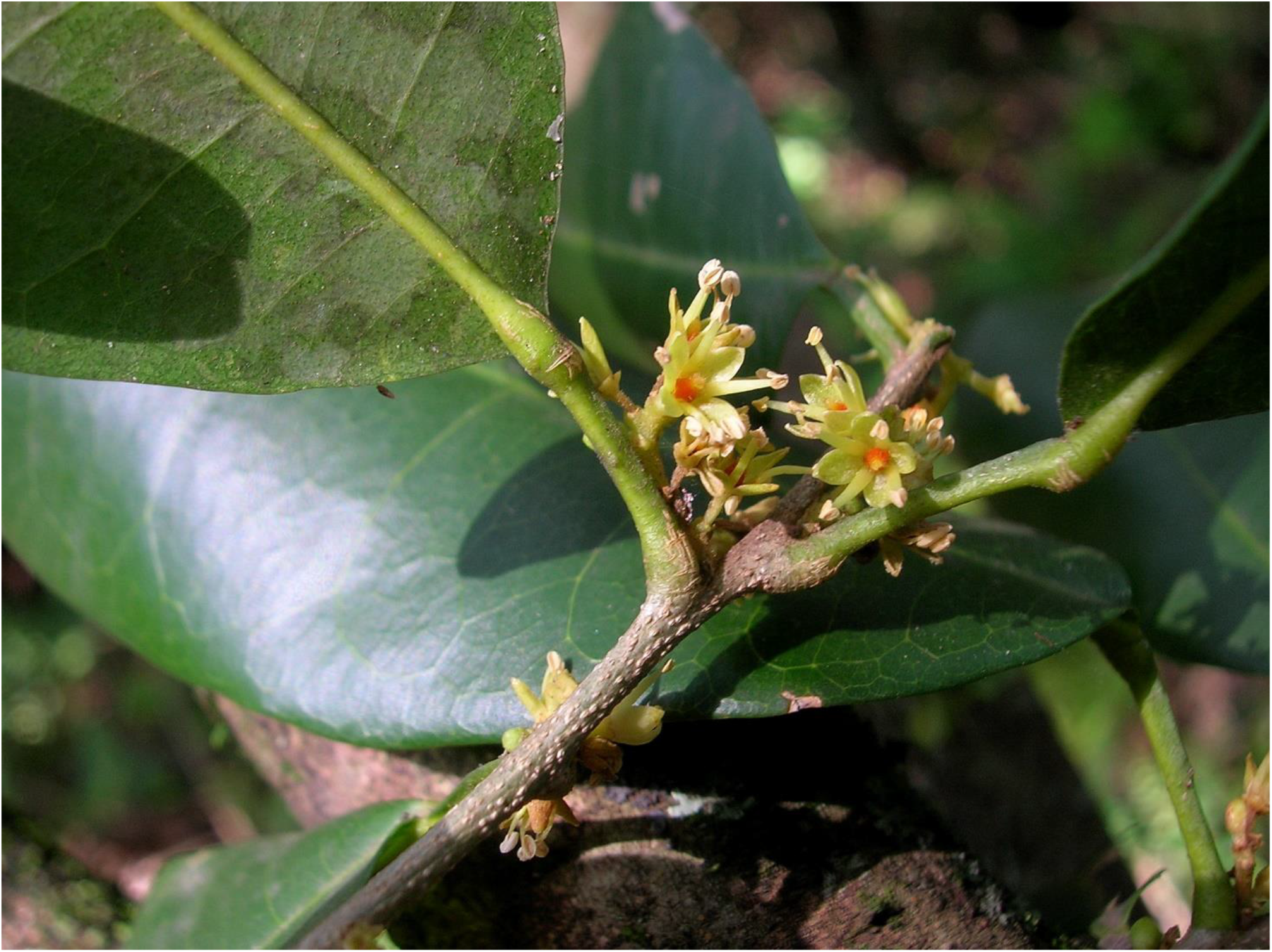
Vepris simplex. Inflorescence with male flowers taken April 2018 on Philip Leakey’s farm in the Loita Hills. Photo W.R.Q. Luke.

Type: Tanzania, “Hochwaldes” (interpreted as Usambara Mts), “1300-1600” m, Sept. 1892, *Holst* 3801 (holotype B, probably destroyed; isotype EA barcode EA000003191!; epitype designated here (see note below) Tanzania, Tanga Province, Lushoto Distr., Manola, 6,600 ft, fl. 16 June 1953, *Parry* 222 (K barcode K000593353!)).

*Vepris simplicifolia* (Engl.) Mziray (Mziray 1992: 75;, White *et al*. 2001: 515) **nom. illegit**.(non *Vepris simplicifolia* Endl. (Endlicher 1833: 89)

*Teclea simplicifolia* (Engl.) I. Verd. (1926: 410; Kokwaro 1982: 25; Gilbert 1989: 427; Friis 1992: 183; Beentje 1994: 369).

*Teclea viridis* I. Verd. (1926: 410). Typ: Kenya, Nairobi Forests, 5500 ft, fl. Feb. 1914, *Battiscombe* 867 (Holotype K 000199480!; isotype EA)

*Teclea unifolioliata sensu* Engl. non Baillon (Engler 1895: 433; 1897: 152)

###### DISTRIBUTION

*Vepris simplex* occurs from the Ethiopian Highlands in the vicinity of Addis Abeba southwards along the E African rift mountains through the highlands of Kenya, and Tanzania (Kokwaro 1982, Friis 1992, Beentje 1994), reaching the Mafinga Mts of northern Malawi (White *et al*. 2001: 515). A putative record from N. Mozambique has not been confirmed by us.

###### REPRESENTATIVE SPECIMENS EXAMINED

**ETHIOPIA.** Mega Mountain, 6300 ft, fr. 9 Sept.,1953, *Bally* 9189 (EA, K!); Sidamo, Mogada, fl. May 1976, *Chaffey* 997 (ETH, K!). **KENYA.** Marsabit, Mt Kulal, 1800 m, fl. Feb. 1959, *T. Adamson* K15(EA, K!); ibid. fr. 29 July 2006, *Nyamongo* in GBK 22 (EA, K!); Kiambu Distr., Nairobi-Nakuru Rd nr Rironi shopping centre, fr 1 Jan. 1976, *Msafiri* 22 (EA, K!); **TANZANIA.** Kilimanjaro, 1800 m, fl. 25 June 1993, *Grimshaw* 93341 (EA, K!); Arusha Distr., Ngongongare forest, fr. 5 May 1960, *Willan* 55 (EA, K!); **HABITAT.** Dry mainly evergreen forest, riverine thicket, evergreen rocky bushland, drier types of upland forest and woodland with *Juniperus* and *Acacia*, extending into the understorey of *Podocarpus* forest (Friis 1992), and in Malawi in montane thicket (White *et al*. 2001). “Common understorey tree at most levels; here in scrub/disturbed relict forest” (*Grimshaw* 94409, K!); 300 – 2300 m altitude.

*Vepris simplex* is by far the most collected species of unifoliolate *Vepris* in tropical Africa, with 317 specimen records on gbif.org. The majority of the specimens were collected in the 1400 – 2300 m altitudinal band in Kenya, extending to the Ethiopian Highlands, and southwards into the high mountains of northern Tanzania: e.g. Kilimanjaro, Mt Hanang, Mt Meru and is morphologically uniform through much of this range, although flowers in Ethiopia are smaller than those in Kenya. In the Arusha area, the leaves are much longer than the norm, oblong and the length: breadth ratio about 3: 1, leaves measuring c. 14.5 × 4.5 cm e.g. *Willan* 514 (K!). In the main part of its range the species often occurs on volcanic rocks such as lava flows and in at least some locations it is “exceedingly common” (Mt Kulal, Kenya, *Bally* 5582, K). Further investigation is needed from specimens from lower altitude evergreen forest areas around Morogoro, Tanzania, and in SE Kenya, e.g. *Magogo & Glover* 693 (Mrima Hill, K!), *Brenan et al*. 14519 (Shaitani Forest near Diani, K!) which are discordant from specimens in the rest of the range. They have large acuminate, papery leaves, exceeding 2:1, with minute, green flowers. These might represent a separate variety or subspecies.

Hermaphrodite flowers with functional ovaries and only two stamens were observed on the otherwise male *Bally* 2578 (K!), by John Hutchinson (specimen annotation).

Trees are predated by elephants *(Grimshaw* 93341, 94409 both K!) which favour this species, and birds eat the fruits *(Grimshaw* 93341, K!), presumably dispersing the seed. This species has the smallest fruits (3 – 5 mm diam.), borne in the greatest numbers per stem than any other unifoliolate *Vepris* species, which may be an adaptation to bird dispersal and contribute to it having the greatest range and being the most frequent of any of the species.

White *et al*. (2001: 516) classify the species as a Sub-Afromontane endemic while Friis (1992) suggests it is an Afromontane endemic (Timberlake 2021c).

*Vepris simplex* is confused with *V. eugeniifolia*. See diagnostic characters under the last species.

###### CONSERVATION STATUS

*Vepris simplex* is listed as Least Concern in view of its vast range and numerous locations, and few specific threats. It is in numerous protected areas in Kenya and Tanzania (Timberlake 2021c).

###### PHENOLOGY

Flowering May-July, fruiting Sept.-Nov. (Ethiopia); flowering June-Feb., fruiting June-Jan. (Kenya), and flowering Nov.-June, fruiting Feb.-June, Aug. (Tanzania).

###### ETYMOLOGY

Originally named simplicifolia by Engler for the unifoliolate (simple not compound) leaves, now known not to be a diagnostic specific character as it must have seemed when first published. Simplex, the new name, coined here, is intended as a convenient, shorter alternative name which is needed since simplicifolia is not available for this species (see Notes, below).

###### VERNACULAR NAMES & USES

Haddessa ormicha (Gallinia, Ethiopia, *Chaffey* 997 (ETH, K!); Mwenderendu (Kikuyu), used for walking sticks (Kenya, *Msafiri* 22 K!); Used for charcoal (Kenya, *Mwangangi* 2344, K!); Goriot (Kips.) and Ol’Gelai (Masai), for walking sticks and bows (Kenya, Narok, *Glover et al*. 22, K!); Kuriot (Kips.) and Olkisi (Masai) for bows, sap for chest troubles (Kenya, Narok, *Glover et al*. 2073, K!); Mulati (Kirangi), used for fuel and building poles (Tanzania, Kondoa Distr., *Ruffo* 781 K!); Engelai (Masai) (Tanzania, *Carmichael* 387, K !); Ndireto (Kimeru) (Tanzania, *Willan* 236, K !); (Ligua) (Tanzania, *Semsei* in FH 2946, K!); Mkuku (Bagamoyo, Tanzania, fide Engler 1895). In addition, the names Muchimi wa Tsakani (Digo), Muretu (Meru), Kurionde(Tugen), Edapalakuyen (Turkana), and the use of wood for roof beams and other artefacts is reported (Beentje 1994).

###### NOTES

The only surviving original material of Engler (1895) located, is the isotype at EA. It is sterile, although the protologue is based on flowering material. Therefore, an epitype is needed to buttress the application of the name to the species and remove ambiguity. There seems some uncertainty about the locality. On the EA isotype label an undecipherable word followed by “Hochwald” (high forest) is written. The location has been inferred or interpreted at a later date by a note in pencil on the label as “Usambara Mts”. However, in the protologue “Usambara Mts” are not mentioned, although an altitudinal range of 1300 – 1600 m is given. The only place name given on the label is Bagamoyo which was the capital of what was then German East Africa and is a historic coastal port town near to the Usambaras. Polhill and Polhill (2015: 199) give an itinerary for Holst in 1892, the year that he collected the original specimen. He was entirely in FTEA T3 (the botanical province containing the Usambara Mts). Given these facts, an epitype has been selected of a fertile specimen representative of the species, also from the T3 area, this is *Parry* 222 (EA, K), chosen because it is of good quality, is in flower showing the representative large male flowers with four stamens, and with the thick, ovate-elliptic leaflets with rounded apices that together unambiguously indicate this species.

When Mziray (1992) made the combination *Vepris simplicifolia*, he was probably unaware that the name was already occupied by *Vepris simplicifolia* Endl. (Endlicher 1833), which is contrary to the Code (Turland *et al*. 2018). The last name was coined for a plant from Norfolk Island in the western Pacific. It is the basionym for *Sarcomelicope simplicifolia* (Endl.) T.G. Hartley (1982: 369) of Australia and New Caledonia which has many local names and likely uses (Hartley 1982).

Therefore, a new name is needed for the African taxon, which is addressed above. The name *Vepris simplex* was selected since it has the advantage of being similar to the name used for the last nearly 100 years, but sufficiently different to be allowed under the Code. It is also less cumbersome, being shorter, always an advantage for users.

Mziray (1992) making the combination *Vepris simplicifolia*, incorrectly gave the authorship as (I.Verd.)Mziray, mistakenly attributing authorship of the basionym to Verdoorn, when she had made it clear that she was making a combination based on Engler’s (1895) *Toddalia simplicifolia*(Verdoorn 1926). Therefore the correct authorship of her name isr *Teclea simplicifolia* (Engl.) I. Verd. and that of Mziray’s is *Vepris simplicifolia* (Engl.) Mziray.

### A note on the Eastern Arc Mountains and Coastal Forests of East Africa

The three new species published in this paper are restricted to the Eastern Arc Mountains and Coastal Forests (EACF) of East Africa, in Tanzania and southern Kenya. The EACF form an archipelago-like phytogeographical unit well-known for high levels of species endemism in many groups of organisms (Gereau *et al*. 2016). Among the better-known mountain blocks are the Nguru Mts, the Udzungwa Mts, the Uluguru Mts, and the Usambara Mts. Supported by moist air currents from the Indian Ocean, the surviving evergreen forests of the Eastern Arc Mountains alone have 223 species of endemic tree (Lovett, 1998), and are variously stated to have 800 (Tanzanian Forest Conservation Group, undated) or as many as 1500 species (Skarbek 2008) of endemic plant species. In herbaceous groups such as the Gesneriaceae, over 50% of the taxa (23 endemic species and a further nine endemic taxa) for East Africa (Uganda, Kenya and Tanzania) are endemic to the Eastern Arc Mts (Darbyshire 2006) and in the Acanthaceae, there are numerous endemic species in multiple genera endemic to the Eastern Arc Mts, e.g. *Stenandrium warneckei, Isoglossa bondwaensis, Isoglossa asystasioides* and *Sclerochiton uluguruensis* (Vollesen 2008; Darbyshire 2009; Darbyshire *et al*. 2010; Darbyshire & Kelbessa 2007). In terms of documented plant species diversity per degree square, the Eastern Arc Mts are second only in tropical Africa to Southwest Cameroon in the Cross-Sanaga Interval of West-Central Africa (Barthlott *et al*. 1996; Cheek *et al*.2001). Several forest genera have disjunct distributions, being found only in the Cross-Sanaga Interval and in the EACF and not in between, e.g. *Zenkerella* Taub. and *Kupea* Cheek (Cheek *et al*. 2003; Cheek 2004c). The EACF include the sole representatives of plant groups otherwise restricted on the continent to the forests of Guineo-Congolian Africa, e.g. *Afrothismia* Schltr. and *Ancistrocladus* Wall. (Cheek & Jannerup 2006; Cheek *et al*. 2000). Extensive forest clearance within the last 100–150 years has removed forest from some mountains entirely, and reduced forest extent greatly in others. Since the 1970s more than 12% of these forests have been cleared (Tanzania Forest Conservation Group, undated). However, forest clearance has appeared to stabilize in the last ten years (Tanzania Forest Conservation Group, undated) in many but not all areas important for plant conservation giving hope that species extinctions can be avoided, or at least kept to a minimum

### Conclusions

The published and provisional extinction risk assessments of the 13 unifoliolate continental *Vepris* species treated in this synopsis indicate that all but two are threatened. Thankfully, the three new species to science published in this paper are all at the lower level of extinction risk, Vulnerable, as a result of the higher levels of protection in the Udzungwa Mountains National Park of Tanzania (*Vepris lukei* and *V. udzungwa*), and the local community protection of the indigenous people of SE Kenya of their Kaya forests (*Vepris roberstsoniae*). However, the future for the three species indicated as Critically Endangered seems fragile or even non-existent. The forest habitat of *Vepris laurifolia* in western Africa (Guinea to Ivory Coast) is steadily being reduced by development projects of multiple sorts including mining and hydropower, and by clearance of the last scraps for agriculture. The two species restricted to the Uluguru mountain forests are of highest concern because since they were last seen nearly 100 years ago, each from a single plant (so far as we are aware) their forest habitat has seen massive clearance. According to Ndang’ang’a *et al*. (2007), the Ulugurus had the highest losses of forest of all Tanzanian EACF areas 1970s-2000 with about 12% loss. All forest is considered to have been lost below 1800 m alt. (East African Plant Red List Authority pers. comm. to first author). It may be that both species are already extinct, in the case of *Vepris* sp. A of FTEA (only recorded below 1800 m alt.), even before it has a scientific name or a formal IUCN conservation assessment published. Until species are scientifically named, it is difficult for an IUCN conservation assessment to be published (Cheek *et al*. 2020b, although there are exceptions, as in *Vepris robertsoniae* of this paper). Most new species to science published today, such as those in this paper, are range-restricted, meaning that they are almost always automatically threatened, although there are exceptions, such as the widespread *Vepris occidentalis* Cheek & Onana (Cheek *et al*. 2019a). Documented extinctions of plant species are increasing (Humphreys *et al*. 2019) and recent estimates suggest that as many as two fifths of the world’s plant species are now threatened with extinction (Nic Lughadha *et al*. 2020). Global extinctions of African plant species continue apace. At the foot of the Udzungwa Mts, the achlorophyllous mycotrophs *Kihansia lovettii* Cheek and *Kupea jonii* Cheek (Triuridaceae, Cheek 2004c) are likely extinct as a result of the placement of the Kihansi hydroelectric dam, not having been seen since construction in 1994 (28 years ago), despite targeted searches. Although not directly threatened by development, another mycotroph, this time in one of the forest fragments of SE Kenya, *Afrothismia baerae* (Thismiaceae, Cheek 2004b) has also not been found despite monitoring in the last 10 years. Global extinctions have also been reported in Guinea, such as *Inversodicraea pygmaea* G.Taylor, and in 2022 after first collection in 2018, *Saxicolella deniseae* Cheek (both Podostemaceae, both extinct due to hydropower construction, Couch *et al*. 2019, Cheek *et al*. 2017; Cheek *et al*. 2022c). New extinctions have recently been reliably reported from Gabon (Moxon-Holt & Cheek 2021, Cheek et al 2021) and Cameroon (Cheek & Williams 1999, Cheek *et al*. 2018c, Cheek *et al*. 2019), including species of *Vepris* (Cheek *et al*. 2018a). If future extinctions are to be avoided, improved conservation prioritisation exercises are needed such as Important Plant Area programmes (Darbyshire *et al*. 2017), supported by greater completion of Red Listing, although this can be slow and problematic (Bachman *et al*. 2019). and, globally, only 21 – 26 % of plant species have conservation assessments (Bachman *et al*. 2018). Where possible, as an insurance policy, seed banking and cultivation of threatened species in dedicated nurseries are urgent. Above all, completion of botanical taxonomic inventories is needed to feed into these exercises, otherwise we will continue to lose species before they are even discovered for science, and certainly before they can be investigated for their potential for beneficial applications. New compounds to science with high potential for humanity are being discovered in *Vepris* species each year (e.g. potent antimicrobial compounds in *Vepris africana*, Langat *et al*. 2021). Such discoveries will not be possible if species extinctions are allowed to continue.

## Acknowledgements

QL would like to thank Tom Butynski and Carolyn Ehardt for allowing him to accompany their primate expeditions in the UMNP funded by Zoo Atlanta and the Margot Marsh Biodiversity Fund We thank Nina Davies for facilitating the loans and working spaces needed for this paper in the Kew Herbarium. Poppy Lawrence assisted in the early stages of this project. Janis Shillito typed the manuscript.

